# *Tubastraea coccinea* (Cnidaria: Anthozoa) is the globally dispersed species of Sun Coral

**DOI:** 10.1101/2025.11.26.690498

**Authors:** Angela Ferreira Portella, João Gabriel Rodinho Nunes Ferreira, Giordano Bruno Soares-Souza, Danielle Amaral, Karine Venegas Macieira, Luciano Cesar Candisani, Yasmin Cunha, Camilla Santos, Daniela Batista, Luciana Leomil, André Torres, Francesco Dondero, Aline Silva Romão-Dumaresq, Diego Figueroa, David Hicks, Juliana Americo, Marcela Uliano-Silva, Claudia A.M. Russo, Mauro de Freitas Rebelo

## Abstract

Species of the genus *Tubastraea* (sun corals) have dispersed globally over recent decades, but morphological trait identification has led to taxonomic confusion, with multiple authors proposing conflicting species identifications due to high morphological plasticity and overlapping diagnostic traits. We combined global-scale molecular phylogenetics with chromosome-level genome assemblies to resolve this taxonomic issue. Using mitochondrial and nuclear markers from 287 specimens across five ocean regions, including type localities in Bora Bora (*T. coccinea*) and Galapagos (*T. tagusensis*), we reconstructed phylogenetic relationships and inferred demographic history through approximate Bayesian computation. We also generated high-quality, chromosome-level genome assemblies for three morphologically distinct morphotypes from the Brazilian coastline. Phylogenetic and species delimitation analyses revealed two well-supported clades: a small, geographically restricted *T. tagusensis* clade confined to the Galapagos Islands, and a large cosmopolitan clade containing all worldwide samples, including *T. coccinea* from its type locality. Chromosome-level genomic analyses confirmed that the three morpho-types represent a single species, sharing a triploid karyotype of 14 chromosomes with extensive synteny and gene-tree discordance consistent with incomplete lineage sorting. Dispersed-route analyses identified the Centralwestern Pacific as the ancestral source, with at least three independent, human-mediated colonization events establishing populations in the Southwestern Atlantic, Gulf of Mexico, and Western Pacific. However, preliminary analysis suggests a Galapagos (Southeastern Pacific) origin rather than Centralwestern Pacific for the *T. tagusensis* and *T. coccinea* cluster. Our results definitively establish *Tubastraea coccinea* as the cosmopolitan species responsible for global dissemination, while *T. tagusensis* is endemic to the Galapagos and not involved in the worldwide dis-persion.

## Introduction

Accurate species delimitation is essential to investigate distribution over geographical space and to identify the evolutionary forces that shape these limits (Ugarte and Garraffoni, 2024). In invertebrates, the task is difficult as these animals often show few diagnostic traits, substantial cryptic diversity and high morphological plasticity. Defining species boundaries in those animals poses a challenge even for the most experienced taxonomists (Knowlton, 1993; Ugarte and Garraffoni, 2024).

Species of the genus *Tubastraea*, commonly known as sun or orange cup coral, exemplify such complex taxonomic issues. Over the past few decades, the geographic range of some species has rapidly expanded to near-cosmopolitan distribution in the Persian Gulf (Hodgson and Carpenter, 1995; Carpenter et al., 1997), the Southwestern Atlantic (de Paula and Creed, 2004; Fenner and Banks, 2004; Bastos et al., 2022; Hoeksema et al., 2024), the Indian, and the eastern and western Pacific (Cairns, 2000).

The fast pace of global expansion for *Tubastraea* clearly points to a human-driven process. Jurisdictional compliance imposes a substantial burden on the maritime sector on account of the sun coral concern (Creed, 2016; Lages et al., 2011; of the Environment – MMA et al., 2018). This includes additional maintenance protocols and periodic shutdowns for vessels and offshore installations, imposing a cumulative cost of hundreds of millions of dollars over the years. Prospective liabilities are expected to escalate due to infrastructure decommissioning, with *Tubastraea* introducing an uncertainty on the order of ten billion dollars in the industry (Guanal-dini, J.H., personal communication, 2022). Nevertheless, taxonomists struggle to determine which sun coral species is responsible for the rapid global colonization. Three species are currently recognized as cosmopolitan (Creed, 2016; Bastos, 2020; Couto et al., 2021; Hoek-sema et al., 2024), *T. micranthus* (Ehrenberg, 1834), *T. coccinea* (Lesson, 1830), and *T. tagusensis* (Wells, 1982). *T. micranthus*, with its black/olive-green corallites, is easily distinguishable from the other two, with bright red-to-yellow corallites. The distinction between *T. coccinea* and *T. tagusensis*, however, has proven much more challenging (Dutra et al., 2023). *T. coccinea* exhibits a color range from bright red to orange corallites (de Paula and Creed, 2004). *T. tagusensis* can also display orange corallites, although, in some localities, yellow coloration is observed (de Paula and Creed, 2004; Sammarco et al., 2010). As corallite coloration ranges overlap and other morphological characteristics fail to distinguish between these species, taxonomic complications arise.

For example, the morphotype commonly identified as *T. tagusensis* in the Southwestern Atlantic by some researchers (de Paula and Creed, 2004; de Oliveira Soares et al., 2016) has been designated as *Tubastraea* sp. cf. *diaphana* by others (Capel et al., 2018), even though the latter has smaller colonies with more polyps, taller and dark brown corallites, as opposed to the bright red-to-orange coloration of *T. tagusensis*. Bastos et al. (2022) suggests that the Southwestern Atlantic populations currently attributed to *T. tagusensis* (orange-yellow corallite) differ morphologically from the Galapagos holotype (bright yellow corallite) and may actually represent a distinct taxon. Therefore, morphological characters alone seems to be insufficient to properly discriminate between these *Tubastraea* sun coral species (Arrigoni et al., 2014; Capel et al., 2017)

Numerous studies have analysed *Tubastraea* corals using molecular data (Capel et al., 2019; Bastos et al., 2022; Costantini et al., 2025), but they lacked the global-scale sampling required to properly determine connectivity in a so-callede cosmopolitan species. In addition, those studies have not sampled the actual types or specimens collected from the type localities, a requirement for the nominal designation of a particular biological species.

In this study, we used molecular markers in a likelihood framework on a global scale to determine the species responsible for the global dispersion of *Tubastraea*. We included samples from type localities: Vaitape Bay (originally Beula Bay) in Bora Bora in French Polynesia in the Indo-Pacific Ocean for *T. coccinea* (Lesson, 1830), and Tagus Cove in the Galapagos archipelago in Ecuador for *T. tagusensis* (Wells, 1982). We also included other samples from the Southeastern, Centralwestern Pacific and the Western Pacific (Indonesia and Vietnam), as well as the Southwestern Atlantic (Brazilian coastline) and the Gulf of Mexico (Bolivar Peninsula). We applied the approximate Bayesian computation to infer the origin (source population) and the routes for the global dissemination.

Moreover, we generated high-quality, chromosome-level genome assemblies for three *Tubastraea* Brazilian morphotypes to examine their morphological differences from a molecular perspective (Brazil, Southwestern Atlantic). These morphotypes have been previously identified as *Tubastraea* cf. *coccinea, T*. cf. *tagusensis* and *Tubastraea* sp. 1 (see Results). These genomes allow direct synteny comparisons and other genomic analyses that provide foundational data for studying the phenotypic plasticity and molecular mechanisms underlying the genus’ dispersal capacity.

## Results

We investigated the taxonomic status of two cosmopolitan, globally dispersed *Tubastraea* species: *T. tagusensis* (e.g., de Paula and Creed (2004); Mantelatto et al. (2011)) and *T. coccinea* (Fenner, 1999; de Paula and Creed, 2004; Sampaio et al., 2012).

To ascertain the correct designation of the nominal species, we included the type localities for both *T. tagusensis* and *T. coccinea* as defined by their original descriptions. Our sampling included 287 specimens from the Gulf of Mexico (Bolivar Peninsula - 22 samples) and the Southeastern Pacific (Galapagos, Ecuador - 31 samples), Centralwestern Pacific (French Polynesia: Moorea - 5 samples; Tahiti - 14 samples; Bora-Bora-29 samples), Western Pacific (Indonesia - 15 samples; Vietnam - 1 sample) and the Southwestern Atlantic (Brazilian coastlines - 170 samples) oceans (Table 1).

**Table 1.**
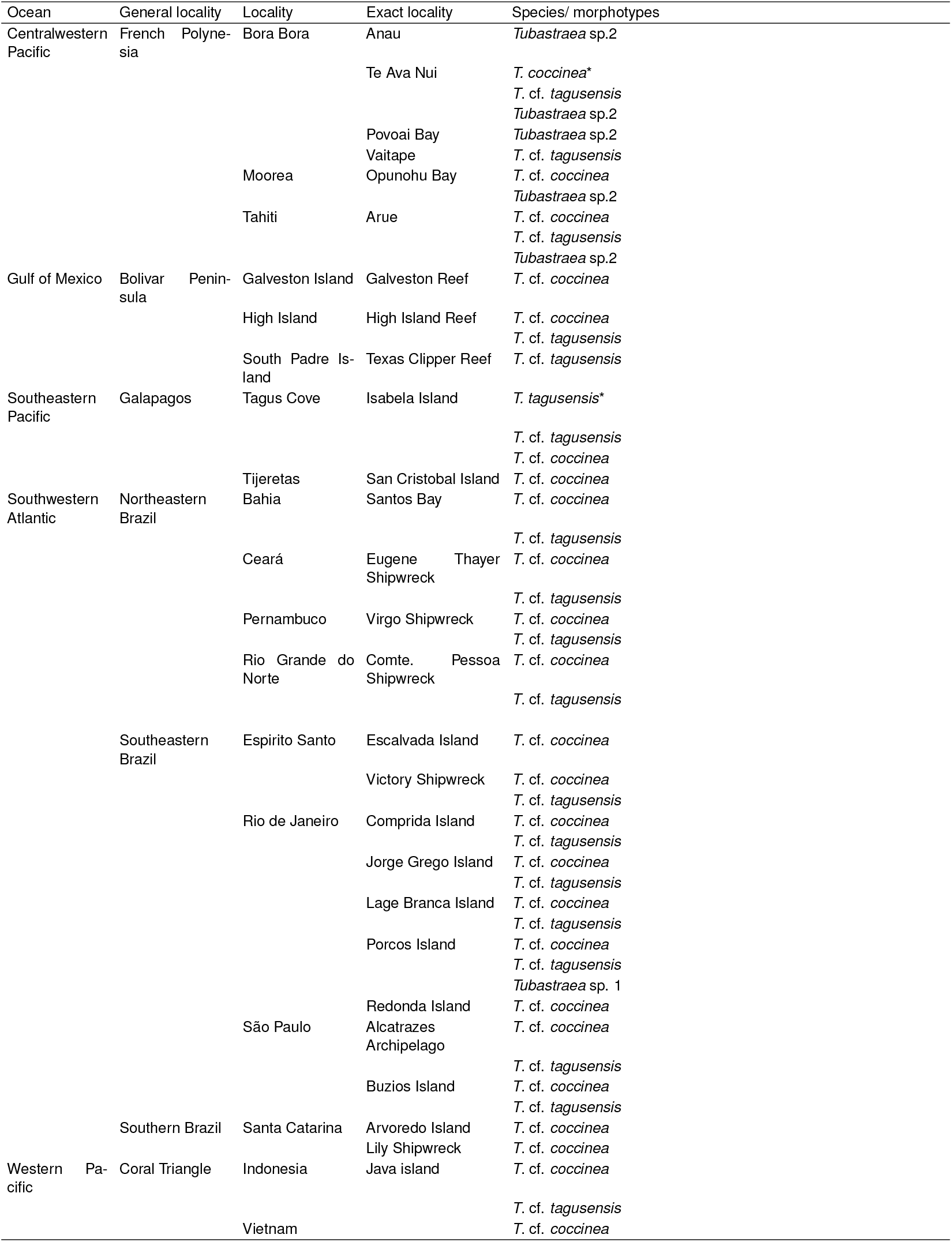
Sampling sites and morphotypes encountered in each location. The asterisks indicate the samples collected on the original locality and used as nominal reference for the species.

The proliferation of distinct *Tubastraea* morphotypes that loosely fit the original descriptions of *T. coccinea* and *T. tagusensis* worldwide (all morphotypes are briefly described in Methods) made sampling challenging. Hence, we restricted the species names for the samples collected in their type localities. In Tagus Cove (Galapagos), for instance, type locality of *T. tagusensis*, we detected three distinct morphotypes. As one of them, detected only in Galapagos, corresponds well to the original and detailed description of *Tubastraea tagusensis* (Wells, 1982), it was designated as *T. tagusensis*. The other morphotype, similar to the one designated as *T. tagusensis* in the Southwestern Atlantic (de Paula and Creed, 2004; Sampaio et al., 2012; Souza Filho et al., 2024), was named *T*. cf. *tagusensis*. The third morphotype is similar to the one assigned to *T. coccinea* worldwide (Sammarco et al., 2014a; Martins et al., 2024; Neves da Rocha et al., 2024) and has been named *T*. cf. *coccinea*.

Similarly, we also detected several morphotypes in Bora Bora, the type locality of *T. coccinea* (Lesson, 1830). Only one morphotype was detected at the entrance of Vaitape Bay, where *T. coccinea* was originally reported. However, this morphotype resembles the aforementioned morphotype *T*. cf. *tagusensis*. In other locations in French Polynesia, we also found this morphotype and, in all cases, it was named *T*. cf. *tagusensis*. The morphotype that corresponds to the original *T. coccinea* description (Lesson, 1830) was found at another location nearby, Te Ava Nui, around 2 miles from Vaitape Bay. Since it complies with the original description and the type locality of *T. coccinea*, it was designated as *T. coccinea*. In the archipelago, we encountered samples of the morphotype that matched the *T*. cf. *tagusensis* from the Southwestern Atlantic and Galapagos and named it *T*. cf. *tagusensis*.

Finally, we also detected other morphotypes that did not fit the ones listed above. Those collected in South-eastern Brazil were designated *Tubastraea* sp. 1 (used in global and genome analyses) and those collected in French Polynesia (used in global analyses)were designated as *Tubastraea* sp. 2. All these designations, used in further analysis, are listed in Table 1.

Our global ML tree (Figure 1) and the ASAP both species delimitation analysis revealed two clades. The first clade includes all seven specimens (SC745, SC766-71) sampled from the type locality of the species in the Galapagos Islands (Tagus Cove, Isabela Island). This clade (marked with black circle; Figure 1) is supported by 100 bootstrap value, a high statistical significance, and includes all samples previously designated as *T. tagusensis*.

**Figure 1.**
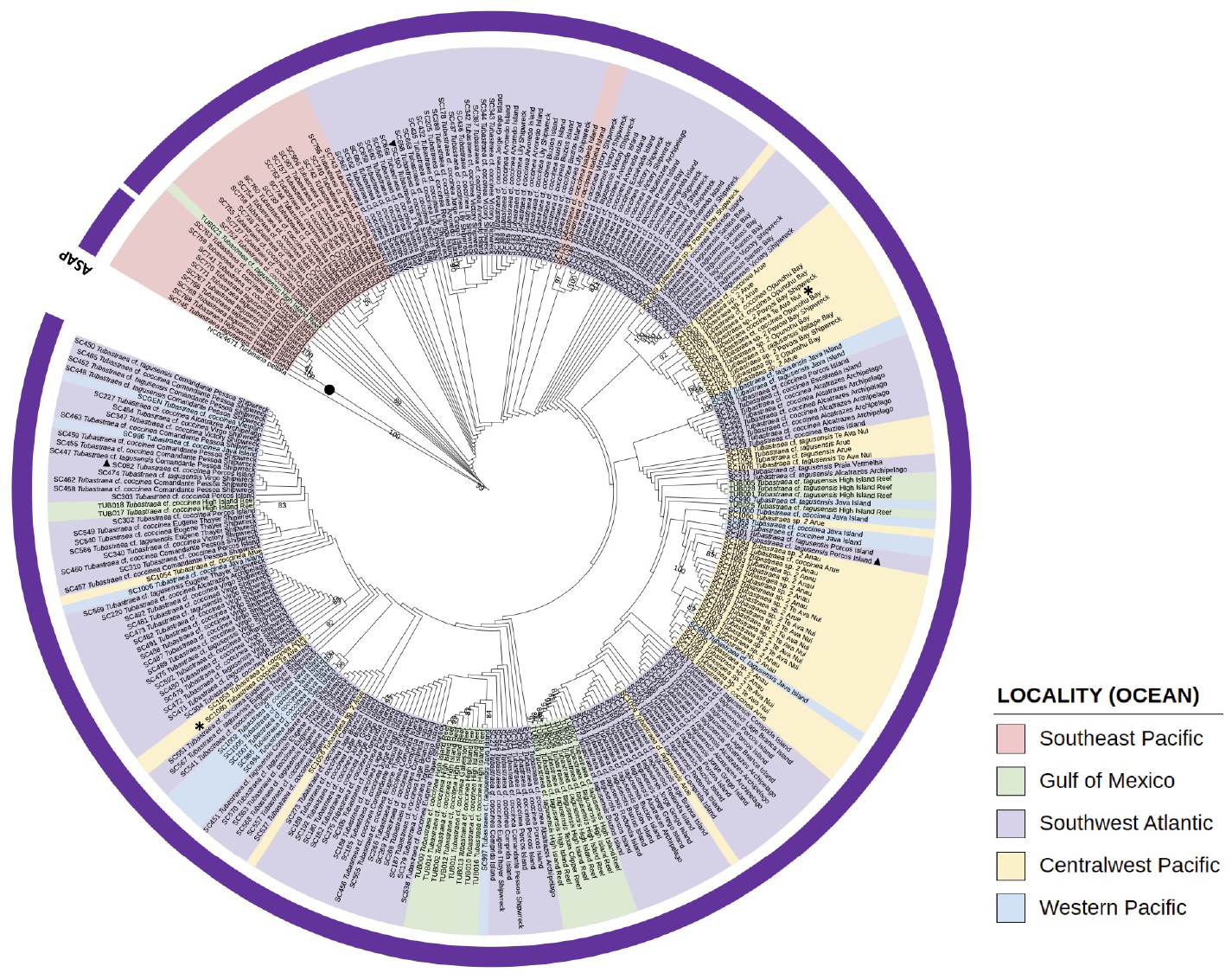
Maximum likelihood phylogenetic tree of *Tubastraea*, based on mitochondrial and nuclear markers. Ultra Fast bootstrap support values are indicated in black. Colored clades correspond to pink = Southeast Pacific; green = Gulf of Mexico; purple = Southwest Atlantic; yellow = Central West Pacific; blue = Western Pacific. See Table 1 for details. Purple circle around the topology delimit species boundaries inferred by ASAP (see Methods for details). The black circle indicates the *T. tagusensis* clade from the Galapagos. The asterisks indicate the two *T. coccinea* sequences from the Bora Bora type locality.

The second clade (Figure 1) includes all worldwide samples identified as *T*. cf. *coccinea*, as *T*. cf. *tagusensis* and as the morphologically variable *Tubastraea* sp. 1 and sp. 2 (bootstrap = 99). It also includes both samples designated as *T. coccinea*, collected at the type locality in Bora Bora (Te Ava Nui). Bootstrap support across the large clade is generally low (<50), forming no distinct clusters distinguishable by morphotype or geography. In our tree, these *T. coccinea* samples appeared in separated clusters (marked with small black stars; Figure 1). Collected in Tahiti, one of them (SC1080) clustered tightly (bootstrap = 92) with a *T*. cf. *coccinea* from Arue (SC1053), also in French Polynesia. The other *T. coccinea* sampled (SC1077) appeared in a low resolution cluster with several *T*. cf. *coccinea, T*. cf. *tagusensis* and *Tubastraea* sp. 2 from French Polynesia and two specimens from Java Island (SC988 and SC989).

We generated chromosome-level reference genomes for the three morphotypes, namely *T*. cf. *coccinea, T*. cf. *tagusensis*, and *Tubastraea* sp. 1, from samples collected in the Southwestern Atlantic in Rio de Janeiro on the Brazilian coastline. Genome profiling with k-mer analysis estimated genome sizes ranging from 527 to 573 Mb (7 Supplementary Figure). Notably, all three morphotypes displayed k-mer spectra indicative of triploidy, a previously unreported feature for the genus (8 Supplementary Figure). Manual curation confirmed a karyotype of 14 chromosomes for all morphotypes (Figure 2; 9 Supplementary Figure). Merqury estimates and BUSCO confirmed low base-level error rates and high completeness (10 Supplementary Figure; 3 Supplementary). Together, these results indicate that the three *Tubastraea* assemblies presented here are high-quality genomic references, providing a robust resource for downstream comparative and evolutionary analyses.

**Figure 2.**
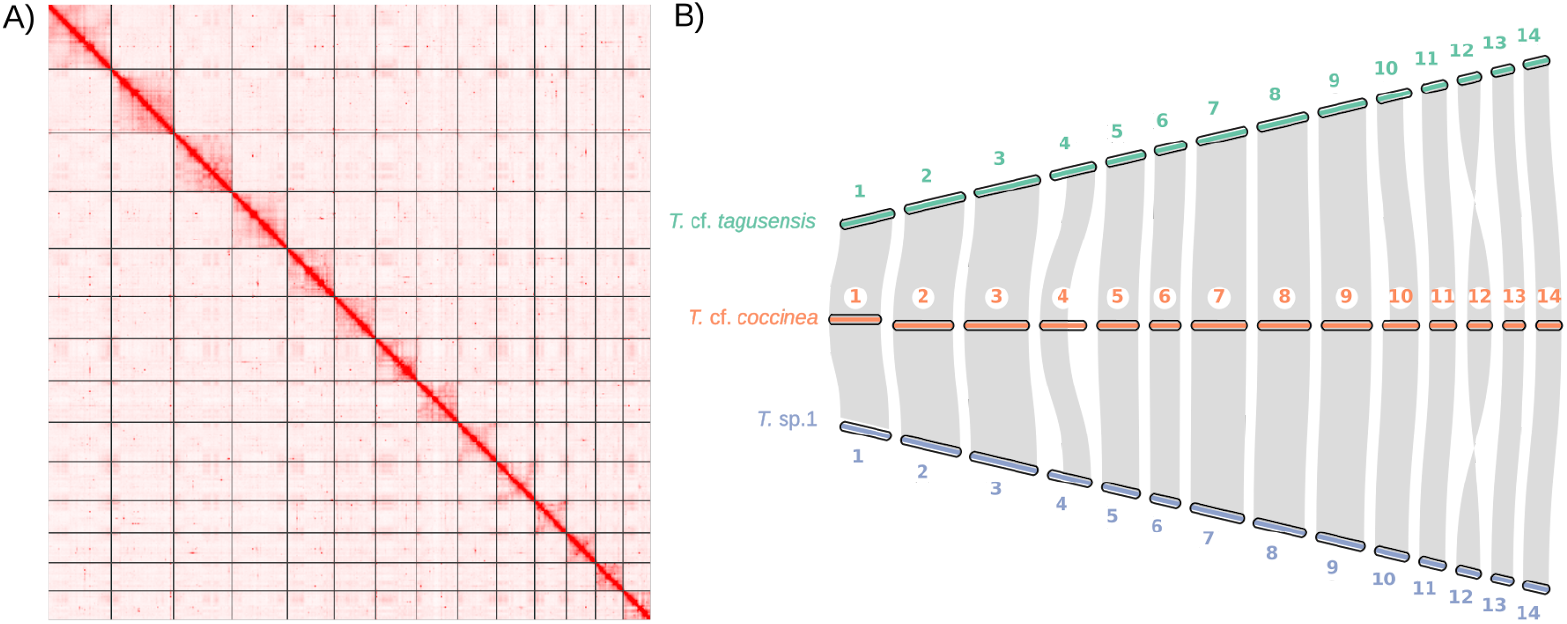
Overview of genome and chromosomal architecture analyses in *Tubastraea*. A) Hi-C contact map of the *T*. cf. *tagusensis* assembly, displaying 14 chromosome-level scaffolds (pseudochromosomes) along the diagonal ordered by decreasing length. (See Supplementary Figure 11 for the other morphotypes). B) A macrosynteny plot showing large-scale gene-order conservation across the 14 reconstructed chromosomes for the three *Tubastraea* morphotypes, based on conserved metazoan BUSCO genes (metazoa_odb10).

Macrosynteny plots based on shared single-copy BUSCO genes showed extensive one-to-one chromosomal correspondence among the three morphotype genomes (Figure 2), with translocations detected in chromosomes 4 and 12 of *T*. cf. *coccinea*, also shown in pairwise whole-genome alignments (11 Supplementary Figure). Altogether, the pronounced collinearity among these genomes supports a shared karyotype among morphotypes, despite their morphological differences in color and shape. To investigate the phylogenetic relationships of those morphotypes, we used 643 singlecopy BUSCO genes predicted for the new chromosome-level genomes presented here. In this analysis, we also included the *T*. cf. *coccinea* genome recently available from Vietnam (Chen et al., 2025) and *Duncanopsammia axifuga* as an outgroup. Both the species tree (coalescent-based ASTRAL) and the maximum-likelihood (ML) tree (concatenated supermatrix) recovered the same topology that places *T*. cf. *tagusensis* as the sister group of the cluster formed by *Tubastraea* sp. 1, *T*. cf. *coccinea* and *T*. cf. *coccinea* Vietnam (Figure 3).

**Figure 3.**
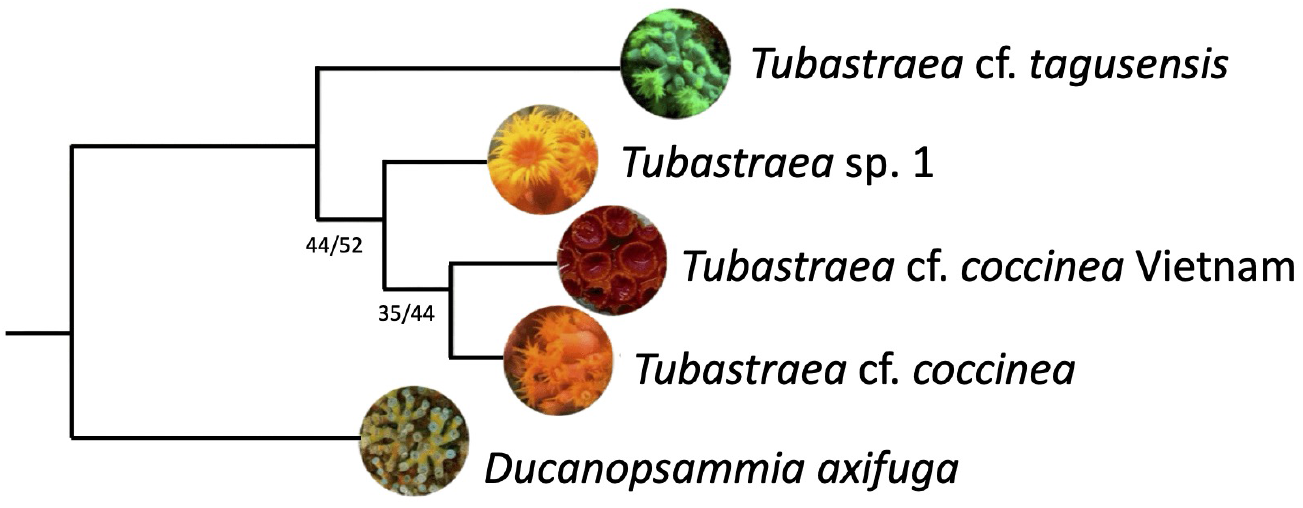
Phylogenomic relationships and gene tree discordance among *Tubastraea* morphotypes. Species tree inferred from the reconciliation of 643 single-copy BUSCO gene trees using ASTRAL-IV, with branch lengths in substitutions per site. All internal branches had maximal support (local posterior probability = 1.0), but LPP values are omitted for clarity. Support values shown at selected nodes represent, respectively, gene concordance factor (gCF) and site concordance factor (sCF). Morphotypes sequenced in this study are shown in bold. *D. axifuga* and *T*. cf. *coccinea* (Vietnam) photographs were sourced from (Subhan et al., 2022) and (Chen et al., 2025), respectively.

Gene and site concordance factors (gCF and sCF see Methods) indicated substantial conflict among loci. Only 44% of gene trees and 52% of informative sites supported the grouping of *Tubastraea* sp. 1 with the two *T*. cf. *coccinea* samples. The closer relationship between *T*. cf. *coccinea* and *T*. cf. *coccinea* Vietnam was supported by 35% of gene trees and 44% of sites (Figure 3). Mapping the genomic positions of discordant genes revealed that these alternative topologies were widely dispersed across chromosomes (12Supplementary Figure). Overall, genome-wide phylogenetic analyses show extensive heterogeneity among loci, mirroring the weak resolution for the large clade observed in the marker-based tree.

## Discussion

### A. Taxonomic status of the cosmopolitan species

Our global-scale analysis used mitochondrial and nuclear markers to determine the taxonomic status of the species responsible for the worldwide expansion (Figure 1). The high statistical support for these clades and the delimitation analysis strongly suggest reproductive isolation between the small *T. tagusensis* clade from Tagus Cove (Galapagos) and the large clade containing all remaining samples, including *T. coccinea* from Te Ava Nui (Bora Bora) (Figure 1).

In the genome-scale analyses, coverage-derived estimates, haplotype structure, and chromosome-scale assemblies are highly conserved (Supplementary Figures 1-4) and genome-wide alignments show conserved synteny across all 14 pseudochromosomes (11 Supplementary). Furthermore, the BUSCO phylogeny and gene-tree results also show no evidence of deep genomic structure separating morphotypes (Figure 3; 12Supplementary Figure). The gene-tree conflict observed, as expected in intraspecific comparisons, reflects incomplete lineage sorting within a single species rather than a historical separation. Despite their distinct appearances, the morphotypes share a triploid karyotype with 14 chromosomes and extensive one-to-one collinearity. Together, the genomic data reinforce the conclusion that the three morphotypes sampled on the Brazilian coastline represent one species.

As the global and genome analyses point to a single biological species, the inclusion of samples collected from the type localities of the species finally settles the taxonomic issue. The cosmopolitan species responsible for the global dissemination is *Tubastraea coccinea*; whereas *T. tagusensis* is confined to the Galapagos Islands and is not a cosmopolitan species. Henceforth, we will keep the name *T. tagusensis* for those samples that appeared in the small clade from Galapagos and samples from the large clade will be referred as *T. coccinea* (including those formerly designated as *T*. cf. *tagusensis, T*. cf. *coccinea, T. coccinea* and *Tubastraea* sp. 1 and *Tubastraea* sp. 2; Figure 1).

The abundance of remarkably variable morphotypes in *T. coccinea* further corroborates the assertion that morphology is a poor predictor of genetic difference in *Tubastraea* (Luz et al., 2020). Phenotypic differences in *T. coccinea* in colony form, corallite density, and even polyp coloration, commonly considered as species indicator (Cairns, 2001; de Paula and Creed, 2004), have already previously been associated with environmental variation such as light availability, depth, and general hydrodynamics (Bastos et al., 2022, 2024). However, the phenotypic nature of this variation could only be confirmed through the morpho-molecular study of the global distribution of *Tubastraea* presented here.

Several new species have recently been described in the literature for the genus, but we found that they require further genetic confirmation. For instance, Yiu and co-workers (Yiu et al., 2021; Yiu and Qiu, 2022) described several new *Tubastraea* species from Hong Kong. However, the phylogenetic trees of those studies indicate that these new species are statistically separated from the *T. coccinea* cluster and are therefore unrelated to the taxonomic issue presented here. In contrast, Serra et al. (2024), based solely on morphology, described species for the Brazilian state of Bahia (Serra et al., 2024). The two morphotypes (*T*. cf. *coccinea* and *T*. cf. *tagusensis*) we sampled in Bahia, however, exhibited no evidence of reproductive isolation from the *T. coccinea* clade. Thus, these new designations must be confirmed with molecular data on a large scale.

Bastos et al. (2022) analyzed several *Tubastraea* morphotypes from Southwest Brazilian and US coastlines. Two of their morphotypes (I and II) match our *T*. cf. *coccinea* and were identified as *T. coccinea*, which agrees with our conclusions. They found a third morphotype (III) in Porcos Island that they designated *Tubastraea* sp., reporting it as genetically distinct from their *T. coccinea* cluster. Their analysis, however, was based on ITS sequences only. Although we also sampled at Porcos Island and collected *T*. cf. *coccinea, T*. cf. *tagusensis*, and *Tubastraea* sp. 1 from this location, we did not encounter the specific morphotype III described by Bastos et al. (2024).

Bastos et al. (2022) designated their third morphotype (III) from Porcos Island as *Tubastraea* sp., reporting it as genetically distinct from *T. coccinea* based on ITS sequences only. However, their analysis was limited to a single nuclear marker (ITS), which may be insufficient for species delimitation given the high morphological and genetic variability we observed within *T. coccinea*. Although we also sampled at Porcos Island and collected *T*. cf. *coccinea, T*. cf. *tagusensis*, and *Tubastraea* sp. 1 from this location, we did not encounter the specific morphotype III described by Bastos et al. (2022). Our genome-scale analyses, using multiple mitochondrial and nuclear markers, showed that morphologically distinct morphotypes (including our *Tubastraea* sp. 1) belong to a single species, *T. coccinea*. Therefore, the genetic distinctness reported by Bastos et al. based on a single marker may reflect intraspecific variation rather than species-level divergence, highlighting the need for integrative approaches combining multiple molecular markers and genomic data for reliable species delimitation in this genus.

A thorough understanding of the morphological variations in *Tubastraea*types will require additional genomic analyses. The translocations on chromosomes 4 and 12 between morphotypes might contribute to such variations, but the available evidence cannot discard epigenetic or developmental mechanisms. These high-quality genomes also enable further investigation focused on genes involved in dispersal and recruitment. Studying reproductive traits and sequencing the geographically restricted *T. tagusensis* from Tagus Cove will be essential to test whether genomic variability explains why *T. coccinea* became cosmopolitan while *T. tagusensis* did not, assuming both experienced similar hydrodynamic and human-mediated opportunities for dispersal.

One key finding in the genomes was triploidy, previously undescribed for this genus. While triploidy has classically been associated with reduced fertility due to meiotic irregularities (Carrasco et al., 1998; Nascimento et al., 2017), triploid organisms can persist through alternative reproductive strategies, including increased reliance on asexual propagation (Vogt et al., 2015). In *Pocillopora acuta* corals, population genomic analyses have revealed a mixed-ploidy system dominated by triploid individuals (*∼*63%), together with strong signatures of predominantly asexual reproduction (Stephens et al., 2023). This pattern may also apply to *T. coccinea*. Capel and co-workers found a high clonality rate (Capel et al., 2019), with more than 70% of their local *T. coccinea* samples sharing identical multilocus genotypes. Although their clonality levels are overestimated, as they treated morphotypes as different species, the clonality levels of *T. coccinea* are likely still high. Rather than limiting the species, triploidy may actually facilitate its global dispersion by promoting asexual reproduction.

### B. Dispersion-route analyses

To understand the dissemination process and possible routes for the cosmopolitan *Tubastraea coccinea* to disperse, we explored alternative scenarios tracing pathways among five geographic populations: the South-western Atlantic (in Brazil: Rio Grande do Norte, Ceará, Pernambuco, Espírito Santo, Rio de Janeiro, São Paulo and Santa Catarina), the Gulf of Mexico (Galveston Island, High Island Reef and South Padre Islands), the Southeastern Pacific (in Galapagos: Tijeretas and Tagus Cove), the Western Pacific (Indonesia and Vietnam), and the Centralwestern Pacific (Moorea, Tahiti, Bora Bora).

Two dispersion-route analyses were conducted. The first (Analysis 1 in Figure 4) focused on determining whether the original source of the *T. coccinea* was the Western Pacific region, the Centralwestern Pacific, or the Southeastern Pacific populations. Our results (PP = 0.468; Figure 4) supported a sequential colonization pattern, with Centralwestern Pacific (Population 1) as the ancestral source, followed by dispersal to the Western Pacific (Population 2) and subsequent expansion to the Southeastern Pacific (Population 3). These results point to a Centralwestern Pacific (native) ancestral source of *T. coccinea*, aligning with early hypotheses of a Centralwestern Pacific origin for the genus (Lesson, 1830; Cairns, 1994).

**Figure 4.**
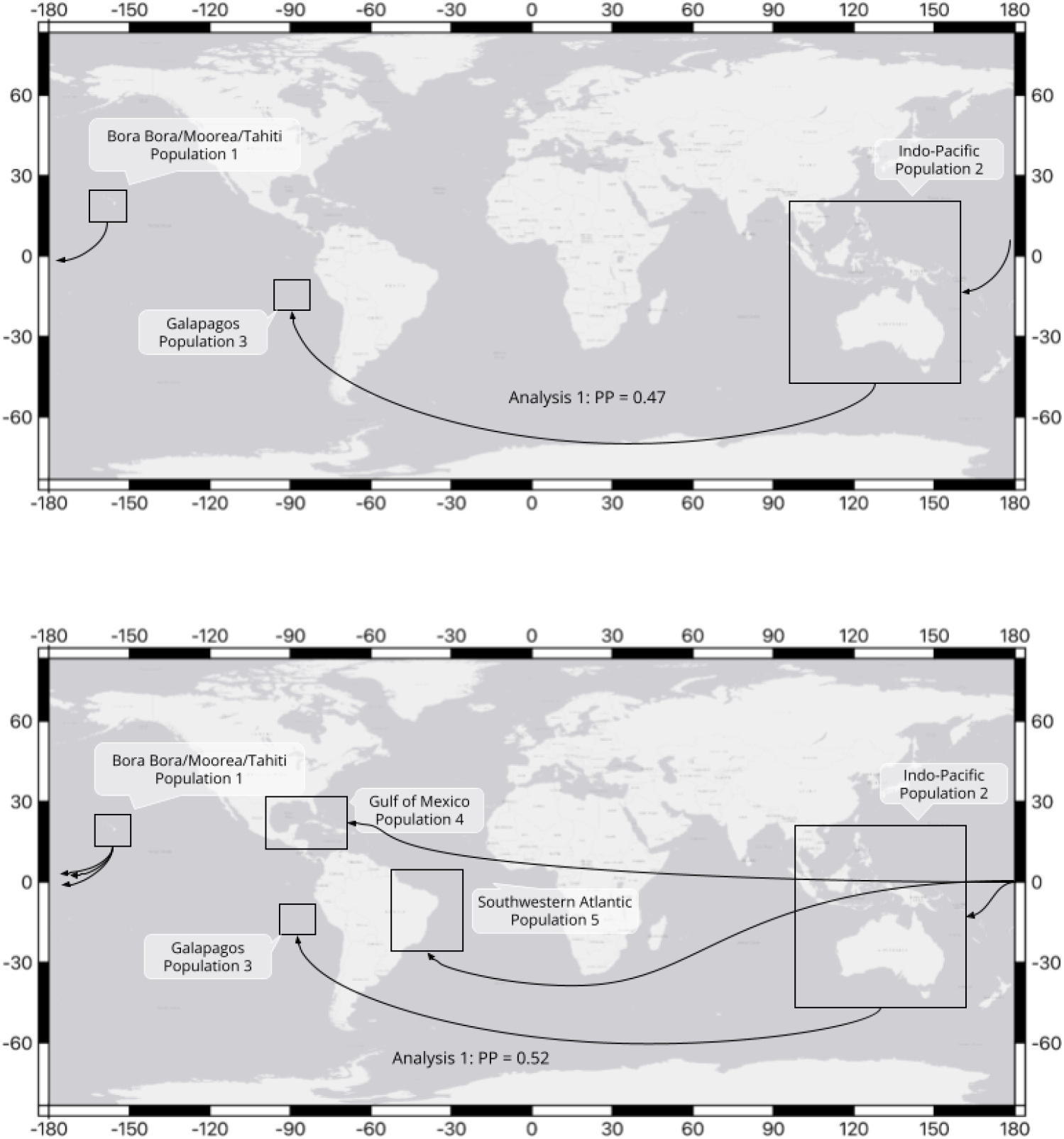
World map showing the sampled populations of *Tubastraea* cf. *coccinea* and DIY-ABC results with COX 1 marker. The populations from the Centralwestern Pacific were considered as a single population, grouped together. This grouped population, along with the Southeastern Pacific population, has been enlarged relative to the map scale for better visualization. Arrows indicate the dispersal routes of the species. (1 - green arrows) Result with the highest posterior probability from the first analysis; (2 - blue arrows) Final result with the highest posterior probability from the second.

On the other hand, early branchings in our global tree (Figure 1) do reveal a clear paraphyletic Galapagos arrangement within the *T. coccinea* clade with most specimens from that locality. This might suggest a Galapagos (Southeastern Pacific) origin rather than Centralwestern Pacific for the *T. tagusensis* and *T. coccinea* cluster. This alternative pattern is the second most supported dispersal route for the origin of *T. coccinea* (Analysis 1). The diversity of *Tubastraea* morphotypes in Galapagos and the difficulty in finding *Tubastraea* in the type location of Bora-Bora (described even by Lesson (1830)) would also indicate a Galapagos origin for the cluster. Confirm this would require a dedicated biogeographical analysis of the species of the *Tubastraea* genus, beyond the aims of this manuscript.

The second analysis (Analysis 2 in Figure 4) explored the demographic connections between Pacific and Atlantic populations, assuming a Centralwestern Pacific origin. It aimed to determine which Pacific population served as the source of the Atlantic colonization, or if multiple events were responsible for the remaining events to the Pacific and Atlantic localities. The second dispersion-route analysis (PP = 0.523; Figure 4) revealed three independent events from the Centralwestern Pacific populations (described below at no particular order).

The first event was responsible for the colonization of the Southwestern Atlantic, encompassing all populations on the Brazilian coastline from Rio Grande do Norte (Northern Brazilian coastline) to Santa Catarina (Southern Brazilian coastline). Interestingly, it seems that *Tubastraea coccinea* arrived in the South Atlantic from the Centralwestern Pacific Ocean (Figure 4), rather than via the Caribbean, despite the shorter distance, high vessel traffic and the early records of the sun coral in the Caribbean in the 1940s (Hoeksema et al., 2024). From the Centralwestern Pacific populations, the second event resulted in the colonization of the Gulf of Mexico (*∼*12,000 km), where populations exhibit low-to-moderate diversity with evidence of admixture among introductions, including the presence of *T. coccinea* and cryptic morphotypes (Figueroa et al., 2019; Bastos, 2020). Secondary dispersal from local hubs such as oil platforms and marinas further homogenizes genetic structure across the Gulf (Sammarco et al., 2012, 2014b). The third event was the colonization of the Western Pacific populations (*∼*7,500 km), including Indonesia and Vietnam.

The Western Pacific emerges as a key redistribution center in the region, bridging the native Pacific with distant regions and facilitating the expansion of the species toward the Southeastern Pacific (Galapagos). The Indonesian port system follows a hub-and-spoke network structure, in which a few highly connected central ports (e.g., Jakarta, Surabaya, Makassar) serve as major nodes linking multiple peripheral sites (Azmi et al., 2015). This configuration facilitates intense inter-island traffic and secondary movements of marine organisms attached to ship hulls, in ballast water, or on transported substrates.

The Western Pacific hosts the world’s most diverse coral reefs, with almost 600 scleractinian coral species across approximately 40,000 km^2^ of reef habitat (Veron et al., 2011; Allen Coral Atlas, 2022). However, rapid degradation due to overfishing, unrestricted tourism, pollution, and climate change (Razak et al., 2022) weakens native community resistance. When combined with dense maritime traffic and frequent handling of coral material, these disturbances provide ideal conditions for opportunistic species such as *Tubastraea coccinea* (Creed, 2016; Soares et al., 2023). Consequently, the Western Pacific’s maritime infrastructure amplifies its ecological role as a secondary dispersal engine, redistributing non-native taxa throughout the region and potentially exporting them to distant oceanic regions (Azmi et al., 2015).As the support was low in the global phylogenetic tree, the number of human-mediated dispersal events cannot be determined (Figure 1).

Similarly, (López et al., 2019) reported thriving *Tubastraea* colonies in the harbor of Santa Cruz de Tenerife (Canary Islands), demonstrating that Atlantic port systems may function as ecological stepping-stones connecting global shipping routes. These findings, along with our data, support the existence of a transoceanic network linking the Pacific, Atlantic, and Mediterranean basins through port-to-port transfers, a pattern consistent with the historical synthesis conducted by Creed (2016).

De Paula et al. (2014) documented several invasion-prone characteristics in *T. coccinea*, including high fecundity (up to*∼* 1,500 eggs per polyp), prolonged reproductive cycles, and larval competency periods sufficient to settle near the parent colony. A high growth rate of up to 1.5 cm per year allows dense colony formation within months, enabling dominance over native sessile fauna (Bastos et al., 2024). These traits ensure rapid recolonization even when transport is sporadic, explaining the swift dispersion observed along the Pacific, the Atlantic, and the Gulf of Mexico in recent decades.

On a molecular scale, Costantini et al. (2025) identified inter-correlated gene modules linked to stress tolerance, mitochondrial performance, and energy metabolism in *T. coccinea*. These gene networks likely underpin their broad ecological amplitude, allowing persistence in turbid, eutrophic, or thermally variable environments typical of harbors. The admixture of divergent lineages can enhance adaptive potential, promote local persistence, and facilitate further expansion. Whether such genetic, ecological and reproductive traits also characterize non-global T. tagusensis remains to be evaluated.

It is also possible that *T. coccinea* has an even more complex history involving intermediate locations from other (unsampled) source populations in the Pacific. The unsampled Florida region, for instance, is widely recognized as one of the major hotspots for biological invasions in the Western Atlantic and a connector between the Gulf of Mexico, the Caribbean, and other tropical areas. South Florida is home to hundreds of established non-native species, driven by intense maritime traffic, high propagule input, favorable climate, and strong habitat modification (Dawson et al., 2017; Searcy et al., 2023). Such conditions suggests that Florida could act as an intermediate point in dispersal routes between the Pacific, the Caribbean, and the Southwest Atlantic. This potential has already been investigated in the Florida Keys and in artificial structures in the Gulf of Mexico, where the sun coral *T. coccinea* has demonstrated the ability to colonize marine platforms and other artificial substrates, forming stable nuclei that can serve as secondary sources of propagules for adjacent natural areas (Precht et al., 2014).

Here we highlight issues arising from reliance on color and visual traits in *T. coccinea*.

We emphasize the importance of sampling original type localities when types are unavailable and an integrative framework, combining global multilocus sampling with genome-scale data. This provides both the geographic breadth and the evolutionary depth required to resolve the taxonomic identity of cosmopolitan species. Together, these findings reveal the interplay between historical biogeographic origins and contemporary human-mediated dispersal in shaping the global distribution of opportunistic marine species such as *Tubastraea coccinea*.

Accurate species identification is therefore essential, as the potential impact of these species on biodiversity hotspots cannot justify hasty reports that lack specialized expertise, which may only exacerbate taxonomic problems (Carlton and Schwindt, 2024).

## Conclusion

We have shown that the cosmopolitan sun/orange cup coral disseminated worldwide is *Tubastraea coccinea*, whereas *T. tagusensis* forms a small, genetically distinct clade restricted to the Galapagos Islands. The extensive morphological variation previously treated as different species or morphotypes represents intraspecific diversity within *T. coccinea*. Genomic analyses of three morphotypes show that they all share a triploid karyotype of 14 chromosomes with extensive synteny, demonstrating that color and shape differences are better explained by phenotypic plasticity rather than genome structure. Gene-tree discordance is consistent with incomplete lineage sorting within a single species rather than structural genomic divergence.

Our dispersion-route analysis identifies the Central-western Pacific as the ancestral source of *T. coccinea*, with at least three independent, human-mediated colonization events establishing populations in the South-western Atlantic, Gulf of Mexico, and Western Pacific. However, there is an indication that the Galapagos may represent the source origin of *T. coccinea*, which would be more biogeographically and genetically plausible, and calls for further analysis to definitively settle this question. These results provide a robust integrative framework for the taxonomy of this genus and highlight how historical biogeography and global maritime traffic jointly shape the distribution of opportunistic marine organisms.

## Materials and Methods

### C. Sample Collection and Morphological Analysis

Colonies of *Tubastraea* were collected by scuba diving on vertical rocky substrates. For global comparative analyses, *T*. cf. *tagusensis, T*. cf. *coccinea*, and *Tubastraea* sp. 1 colonies were collected from seven regions across the Atlantic and Pacific Oceans. Detailed sampling information is provided in Table 1 (see also Figure 5). For the genome analysis, the same protocol was used on Porcos Pequena Island (23°3.229’S; 44°19.058’W) for *T*. cf. *tagusensis* and *T*. cf. *coccinea*, while *Tubastraea* sp. 1 was collected off Porcos Island (22°57’56”S; 41°59’36”W) on the Brazilian coastline.

**Figure 5.**
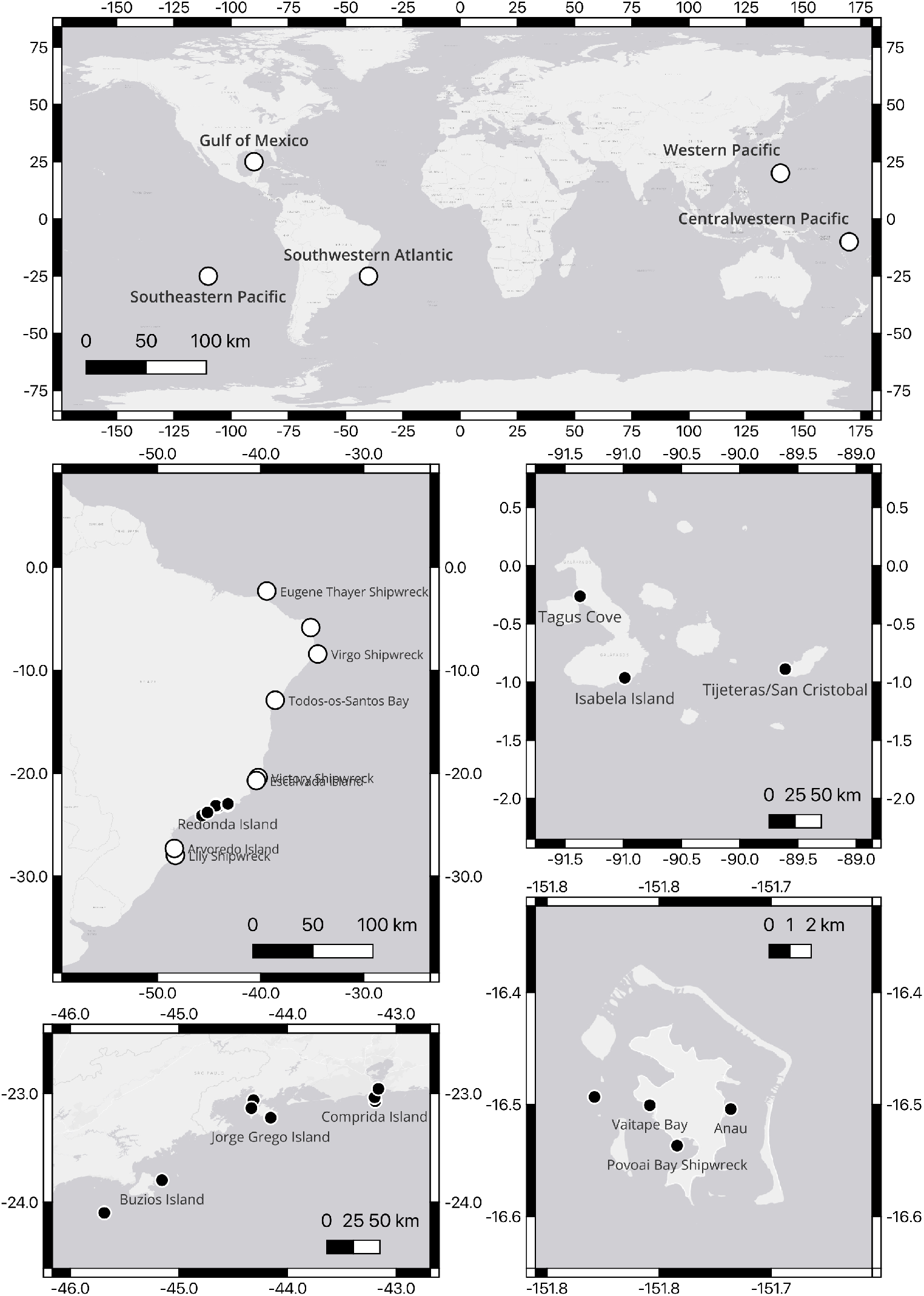
Global and detailed maps showing sampling sites for *Tubastraea* species

For morphological analyses, colonies were immersed in NaOCl for three days to remove soft tissues, rinsed, and air-dried and photographed with a Nikon D750 and Leica M205 FA stereomicroscope. Measurements of colony size, calice diameters and polyp height were taken with calipers. Five distinct morphotypes were recognized and are described below (Figure 6).

**Figure 6.**
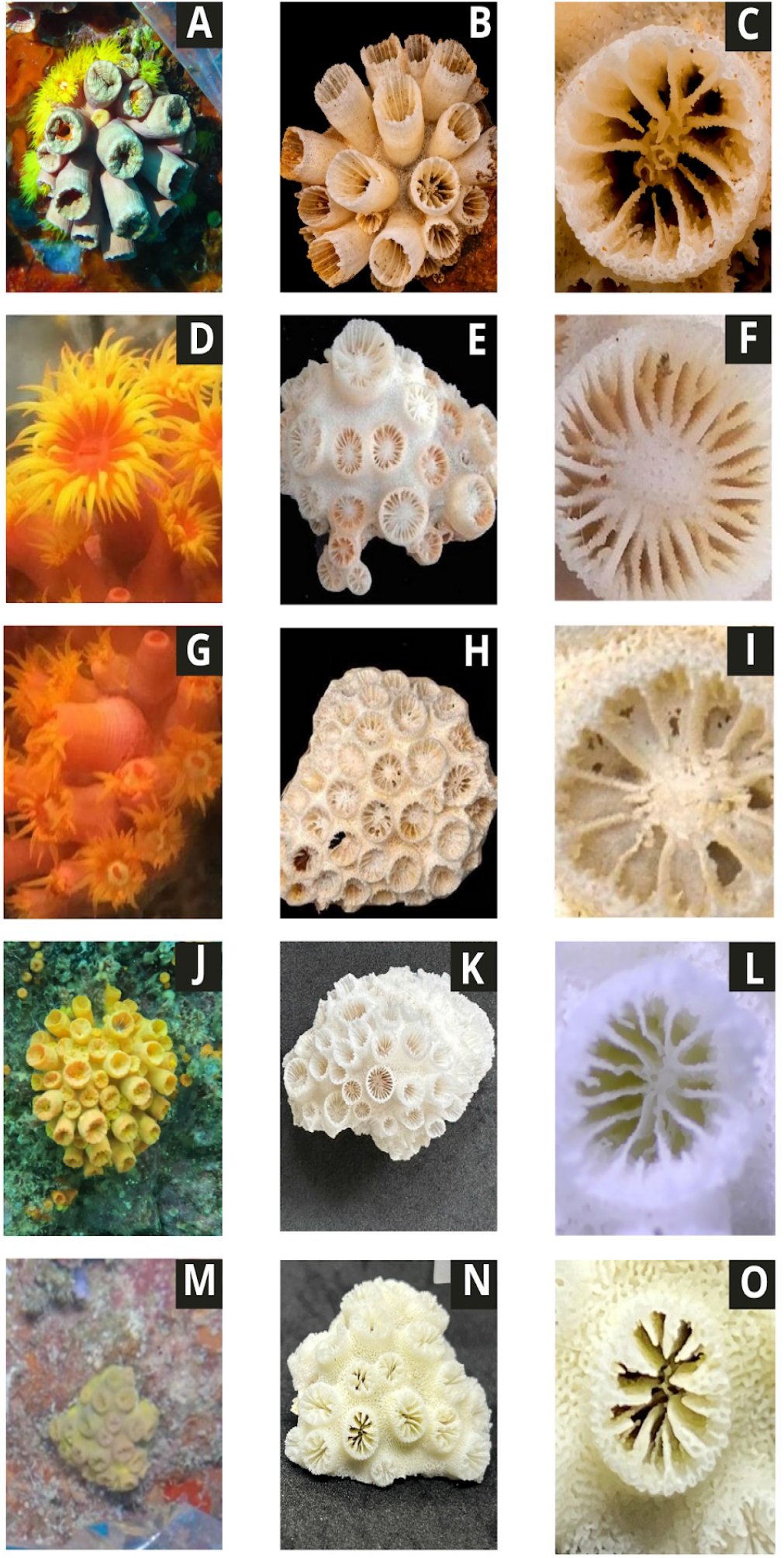
Tubastraea morphotypes analyzed in this study. (A-C) Morphotype of *T*. cf. *tagusensis* (genome analysis, SC450) from Brazil (A) in vivo, with yellow-orange polyps connected by coenosarc with the same color; (B) Corallum phaceloid forming colony with 4.9 cm in diameter; (C) Detail of septa arrangement, with primaries (S1) and secondaries (S2) septa reaching the center of corallite. (D-F) Morphotype of *Tubastraea* sp. 1 (genome analysis, SC100) from Brazil (D) in vivo, used in genome analysis with coenosarc orange-red in color, while tentacles and mouth are yellow and red-orange bright, respectively; (E) Corallum placoid forming colony with 6.7 cm in diameter; (F) Detail of septa arrangement, with part of cycle septal with the initial stage of Pourtalés Plan. (G-I) Morphotype of *T*. cf. *coccinea* (genome analysis, SC082) from Brazil (G) in vivo; (H) Corallum placoid forming colony with 6.7 cm in diameter; (I) Detail of septa arrangement, with S1 and S2 reaching the center of the corallite. (J-L) Morphotype of *T. tagusensis* (SC897) from Galapagos (J) in vivo with bright yellow polyps connected by coenosarc with the same colour; (K) Corallum phaceloid forming colony with 5 cm in diameter; (L) Detail of septa arrangement with primary (S1) and secondary (S2) septa reaching the center of the corallite, whereas tertiary septa (S3) only occasionally do. Corallite with 0.5 cm of diameter. (M-O) Morphotype of *T. coccinea* (SC1077) from Bora Bora (M) in vivo; (N) Corallum placoid forming colony with 3.0 cm in diameter; (O) Detail of septa arrangement, with S1 and S2 reaching the center of the corallite. Row J-L represents *T. tagusensis* and all other rows represent the same species *T. coccinea*

*T. tagusensis* (From the type locality Galapagos Archipelago): Colonies display bright yellow polyps connected by coenosarc of similar color, forming phaceloid colonies up to 8 cm in diameter. Corallites are cylindrical, thin-walled, and loosely arranged (up to 1.0 cm in diameter), with primary (S1) and secondary (S2) septa extending to the calice center, and tertiary (S3) septa occasionally present. These traits closely match the description by Wells (1982) for *T. tagusensis* from Tagus Cove, characterized by small, globular colonies (7-10 cm) of thin-walled, cylindrical corallites (0.8-1.0 cm) with nearly smooth or sparsely granulose walls, low costae, and a finely flaky, non-costate coenosteum.

*T*. cf. *tagusensis* (worldwide, including Galapagos and Bora Bora): Colonies exhibit yellow-orange polyps and phaceloid growth forms reaching up to 13 cm in diameter. Corallites are cylindrical with well-developed septa; both S1 and S2 extend to the center of the calice.

*T. coccinea* (Bora Bora) and *T*. cf. *coccinea* (worldwide, including Galapagos and Bora Bora): Colonies are orange to red with a placoid growth form, approximately 6-7 cm in diameter. Corallites are densely packed, with thick walls, reduced coenosteum, and S1 and S2 septa reaching the calice center. This morphology corresponds to *T. coccinea* as originally described by (Lesson, 1830) from Bora Bora.

*Tubastraea* sp. 1 (Brazil): Colonies display orange-red coenosarc with bright yellow to red-orange tentacles and oral discs. The corallum is placoid, with corallites originating from a shared basal coenosteum and moderately spaced by budding. Mature corallites project up to 10 mm above the coenosteum and exhibit partial septal cycles.

*Tubastraea* sp.2 (French Polynesia): Samples from Bora Bora include more than one distinct morphotype, suggesting that the taxonomic boundaries of *T. coccinea* in this region may encompass multiple forms. Specimens that do not conform to the above descriptions were provisionally referred to as *Tubastraea* sp.2 pending further analysis.

### D. Genome Sequences and Analyses

Coral soft tissue was rinsed with distilled water before nucleic acid extraction. DNA from *T*. cf. *tagusensis* (Illumina) was extracted with the DNeasy Blood & Tissue Kit (Qiagen), while DNA from other samples (*T*. cf. *tagusensis* PacBio, *T*. cf. *coccinea* and *Tubastraea* sp. 1 Illumina and PacBio) followed the CTAB-based protocol of (Garcia et al., 2013). RNA was extracted using the TRIzol protocol (De Paula et al., 2014). Nucleic acid quality was assessed by spectrophotometry, fluorometry, and agarose gel electrophoresis.

High-quality samples (DNA 10-60 kb; RNA RIN *≥*6.6) were used for Illumina library prep (150 bp paired-end, HiSeq X) and NEBNext mRNA libraries. Additional tissue (25 mg) was used for extraction with the DNeasy kit for marker amplification. DNA integrity was checked by agarose gel electrophoresis, PCRs were performed with DreamTaq Master Mix (Thermo Fisher), and purified amplicons were sequenced at Macrogen. Primer pairs used for marker amplification are listed (Table 2). All RNA libraries were prepared with the NEBNext Ultra II RNA Library Prep Kit and Poly(A) mRNA Magnetic Isolation Module (New England Biolabs), following manufacturer instructions. Libraries were pooled and sequenced on an Illumina HiSeq X (150 bp paired-end,*∼*30 M reads per sample). Genomic DNA libraries were prepared with the Kapa Hyper Kit for *T*. cf. *tagusensis* (insert*∼* 494 bp) and the Nextera DNA Flex Kit for *T*. cf. *coccinea* and *Tubastraea* sp. 1 (average inserts 557 and 496 bp, respectively). Sequencing was performed on HiSeq X and NextSeq 550 platforms (Illumina).

**Table 2.**
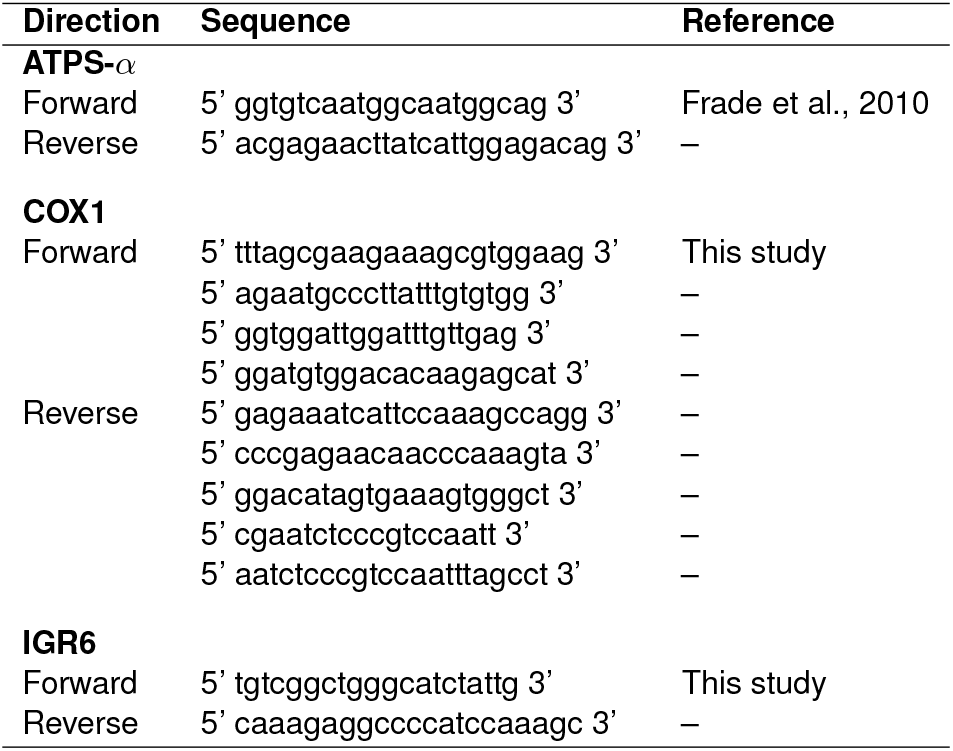
Primer pairs used for PCR amplification of genetic markers in *Tubastraea*.

PacBio SMRTbell libraries for all morphotypes followed the standard PacBio protocol. High-molecular-weight DNA was used directly for *T*. cf. *tagusensis* (*∼* 20 kb) and *Tubastraea* sp. 1 (*∼* 14 kb), while *T*. cf. *coccinea* DNA was sheared with a Covaris g-TUBE. Libraries were size-selected with BluePippin (>15 kb) or AMPure beads, and sequenced on a PacBio Sequel system. Raw Illumina reads from *T*. cf. *coccinea, T*. cf. *tagusensis*, and *Tubastraea* sp. 1 were analyzed with GenomeScope 2.0 to estimate genome size, heterozygosity, and repeat content, and with Smudgeplot v0.2.5 to infer ploidy (Ranallo-Benavidez et al., 2020).

Genome assemblies used MaSuRCA v.3.1.1 (Maryland Super-Read Celera Assembler) (Zimin et al., 2013) with Flye with default parameters apart from JF_SIZE = 30,000,000,000, removing redundant haplotypes with Purge Haplotigs v1.0.4 (Capel et al., 2019). Chromosome scaffolding involved Dovetail Chicago and Hi-C libraries with HiRise and manual curation following the GRIT protocol (Howe et al., 2021). Assembly metrics used the asmstats script, completeness used BUSCO v5.8.2 (metazoa_odb10, genome mode, MetaEuk; (Simão et al., 2015) and base-level accuracy used Merqury v1.1, with k-mer databases (k=21) by meryl v1.3 from Illumina reads (Rhie et al., 2020) (Table 3 for assembly stats; see Data Availability section for accession codes).

**Table 3.** Assembly statistics for the three *Tubastraea* morphotypes.

| Metric | *T. cf. coccinea* | *T. sp. 1* | *T. cf. tagusensis* |
| --- | --- | --- | --- |
| Total assembly length (Mb) | 787.73 | 772.04 | 680.50 |
| GC content (%) | 38.60 | 38.80 | 38.80 |
| Mitogenome length (bp) | 19 092 | 19 092 | 19 093 |
| Number of scaffolds | 4 207 | 2 301 | 622 |
| Number of chromosome-level scaffolds | 14 | 14 | 14 |
| Scaffold N50 (Mb) | 41.28 | 47.98 | 44.34 |
| Scaffold L50 | 8 | 7 | 6 |
| Number of contigs | 17 746 | 13 275 | 6 837 |
| Largest contig (Kb) | 478.86 | 465.79 | 1 261.51 |
| Mean contig (Kb) | 44.31 | 58.07 | 99.41 |
| Contig N50 (Kb) | 61.36 | 76.34 | 156.48 |
| Contig L50 | 4 016 | 3 258 | 1 315 |
| Assembly in pseudochromosomes (%) | 76.14 | 84.42 | 95.24 |
| QV (Merqury, Illumina reads) | 39.23 | 37.04 | 40.13 |
| Reads mapped back (%; Illumina) | 96.54 | 97.44 | 97.18 |
| BUSCO 5.8.2<br>(metazoa_odb10) | C:95.5% [S:91.4%, D:4.1%]<br>F:2.2%, M:2.3%, n:954 | C:91.6% [S:87.7%, D:3.9%]<br>F:3.0%, M:5.3%, n:954 | C:92.8% [S:87.8%, D:4.9%]<br>F:3.0%, M:4.2%, n:954 |

Pairwise alignments among *Tubastraea* genomes were performed using minimap2 (Li, 2018), and dot plots were generated with D-GENIES (Cabanettes and Klopp 2018). BUSCO genes identified with the metazoa_-odb10 dataset were used for both synteny and phylogenomic analyses. Macrosynteny among the three genomes was assessed using the MCscan module of the JCVI toolkit v1.5.4 (Tang et al., 2024), following developer recommendations. Phylogenomic analyses were based on single-copy BUSCOs shared across five scleractinian genomes — the three *Tubastraea* morpho-types from this study, a published *T. coccinea* genome from Vietnam (Chen et al., 2025), and *Duncanopsammia axifuga* as outgroup. Protein sequences were aligned with FSA v1.15.9, codon alignments generated with PAL2NAL v14, and poorly aligned regions removed using GBlocks v0.91b (-t=c -b5=h). Gene trees were inferred individually with IQ-TREE2 v2.4.0 (-m MFP).

Two complementary species tree approaches were then applied. The first was the supermatrix, where individual gene alignments were concatenated with AMAS (Borowiec, 2016). A maximum likelihood tree inferred with IQ-TREE2 (-m MFP -bb 1000 -alrt 1000 -p partition.nex). Gene and site concordance factors were computed (–gcf, –scf 100). The second was the species tree, where individual gene trees were analyzed in ASTRAL-IV (Zhang and Mirarab, 2022), using the CASTLES-II method (Tabatabaee et al., 2023) to estimate branch lengths. Concordance factors were also computed in IQ-TREE2. To visualize phylogenetic signal heterogeneity, a custom Python script (plot_topologies.py) summarized gene-tree topologies using Biopython and Matplotlib, showing the frequency of alternative resolutions among loci.

### E. Molecular Analyses in a Global Scale

For the molecular global analyses, three genetic markers were analyzed to assess genetic diversity and species boundaries: two mitochondrial regions (COX1 and IGS6, the intergenic spacer between ND4 and rns (Capel et al., 2016), and one nuclear gene (ATPS*α*). Although COX1 is widely used for DNA barcoding, it shows limited variability in Anthozoans (Shearer et al., 2002; McFADDEN et al., 2011; Quattrini et al., 2023) and the more variable IGS6 and ATPS*α* markers were also included in our analyses (see Data Availability section for accession codes).

Gene sequences were aligned individually in MAFFT (Katoh et al., 2019). For COX1, the E-INS-i strategy was applied due to the marker’s structural complexity, but, for the IGS6 and ATPS*α*, the faster G-INS-i strategy was used. Alignments were subsequently checked manually and concatenated into a single dataset in MEGA v.11 (Tamura et al., 2021).

As evolutionary rates vary among markers, partitioning was applied before phylogenetic inference (Marshall et al., 2006; Brown and Lemmon, 2007; Kainer and Lanfear, 2015). The best-fit models were selected with ModelFinder (Kalyaanamoorthy et al., 2017) in IQ-TREE2 (Minh et al., 2020): HKY+F+I+R2 for COX1, TIM+F+R2 for IGS6, and K3Pu+I for ATPS*α*. Maximum likelihood analyses were run in IQ-TREE2 with 1,000 bootstrap pseudo-replicates to assess branch support (Felsenstein, 1985; Romano and Cairns, 2000; Holmes, 2003; Takata et al., 2025).

Our phylogenetic inference (Figure 1) was complemented with a species delimitation and a global dispersion-routes analyses (Mera-Rodríguez et al., 2025). For the species delimitation analysis, we used the Assemble Species by Automatic Partitioning (ASAP; Puillandre et al. (2020)). It provides a distance-based method, clustering sequences into species partitions ranked by a scoring system that accounts for barcode gaps and probabilities of panmixia (Puillandre et al., 2020). It is more computationally efficient and does not require prior assumptions about species limits, making it suitable for large DNA barcoding datasets. Genetic distances were calculated using the Kimura 2-Parameter model ((Kimura, 1980)).

For the global dispersion route analysis (Figure 4), demographic scenarios were inferred for five geographic populations of *Tubastraea coccinea*: Southwestern Atlantic, Gulf of Mexico, Southeastern Pacific, Western Pacific, and Centralwestern Pacific. Analyses were based on the mitochondrial marker COX1, chosen for its broad application in marine studies and for enabling comparisons across datasets while remaining computationally efficient (Capel et al., 2016; Seiblitz et al., 2022).

Two analyses were conducted: (1) dispersal among Pacific populations and (2) connectivity between Pacific and Atlantic groups. Alternative scenarios were developed following (Fraimout et al., 2017) and tested independently in DIYABC-RF (Cornuet et al., 2014) with 5 million simulations per analysis. Parameters included Ne = 100,000-1,000,000, divergence times = 10-1,000 generations (Boavida et al., 2019), and mutation rates = 10^*−*7^–10^*−*9^, with r1 = 0.01-0.999. All simulations employed the Kimura 2-Parameter model. Prior distributions were checked through exploratory runs, and the Random Forest module with 1,000 trees was used for model selection and parameter estimation, enhancing discrimination among scenarios (Cornuet et al., 2014).

## Data Availability

All datasets generated in this study have been submitted to the European Nucleotide Archive (ENA) and will be released upon publication. Molecular marker sequences are deposited under OZ364384-OZ364632 (ATPS-*α*), OZ364119-OZ364383 (IGS6), and OZ363218-OZ363478 (COX1).

Raw sequencing data used for genome assembly are available under the following ENA accessions: PacBio CLR WGS (ERR15536567-ERR15536593), Illumina WGS (ERR15535356, ERR15535437, ERR15536492, ERR15536550-ERR15536552), Illumina RNA-seq (ERR15536612-ERR15536614), and Illumina Hi-C (ERR15535066, ERR15535086, ERR15535089). Based on our results, the three assemblies correspond to colonies of *Tubastraea coccinea* and were therefore submitted under this species name: TubCoc1 (*T. coccinea*; GCA_977035605), TubCoc2 (formerly *Tubastraea* sp.; GCA_977035615), and TubCoc3 (formerly *T*. cf. *tagusensis*; GCA_977035595).

## Acknowledgements

Authors thank Dr. Bárbara Segal e Marcelo Crivella from Federal University of Santa Catarina for samples and sharing their knowledge and experience with the *Tubastraea* infestation. We also thank Dr. Emiliano Calderon for providing samples from Macae, Dr. Guilherme Longo from the Federal University of Rio Grande do Norte for samples and Dr. Andrew Calcino for suggestions in the early stage of the work. We want to acknowledge the support of Dr. Margarita Brandt and Dr. Diana Pazmiño from Universidad San Francisco de Quito during the Galapagos expedition, as well as Dr. Benoit Espiau, Dr. Yannick Chancerelle and Dr. Laetitia Hédouin from CRIOBE for their support during the French Polynesian expedition. AI-based tools were used to improve readability and revise the language, not to include, alter or interpret scientific content. This study was funded by Repsol SINOPEC Brazil under Brazilian Petroleum Agency grants 23251-2/2022 and 20771-2/2018. This study was also financially supported by the National Research and Technology Council (CNPq - 315443/2021-9,301659/2025-7) and from Rio de Janeiro State Research Funding Agency (FAPERJ - E-26/010.001887/2019, SEI-260003/001170/2020, SEI-260003/012995/2021, SEI-260003/006126/2024) grants received by CAMR. The funders had no role in study design, data collection and analysis, decision to publish, or preparation of the manuscript.

## Sampling Permits

The collections conducted in this study complied with all legal regulations and were carried out under the appropriate permits issued by each country. Sampling in French Polynesia (Bora Bora, Moorea, Tahaa, and Tahiti) was authorized by the Government of French Polynesia through Arrêté n°ENV24502146AM-1, which permitted the collection and access to genetic resources of *Tubastraea coccinea* and *T. tagusensis* between 18 March and 8 April 2024, as well as the export of material for laboratory analyses. Collections in the Galapagos Islands were conducted in accordance with the Genetic Resources Access Framework (Contract MAE-DNB-CM-2021-0174) and were supported by the corresponding sample mobilization authorization issued by the Agencia de Bioseguridad y Cuarentena para Galapagos, allowing the transport of tissues and colonies of *T. tagusensis* from Isla Isabela to San Cristóbal between November and December 2023. In Brazil, field activities were conducted under SISBIO Authorization No. 86039-5 (ICMBio/MMA), granted to the project “Monitoring of the Sun-Coral (*Tubastraea* spp.) along the Brazilian coast using environmental DNA methodology,” which permits the collection, transport, and temporary maintenance of *Tubastraea* specimens across multiple coastal localities between October 2022 and October 2024. All activities fully complied with environmental regulations and legal requirements of each jurisdiction.

## Supplementary Materials

**Figure 7.**
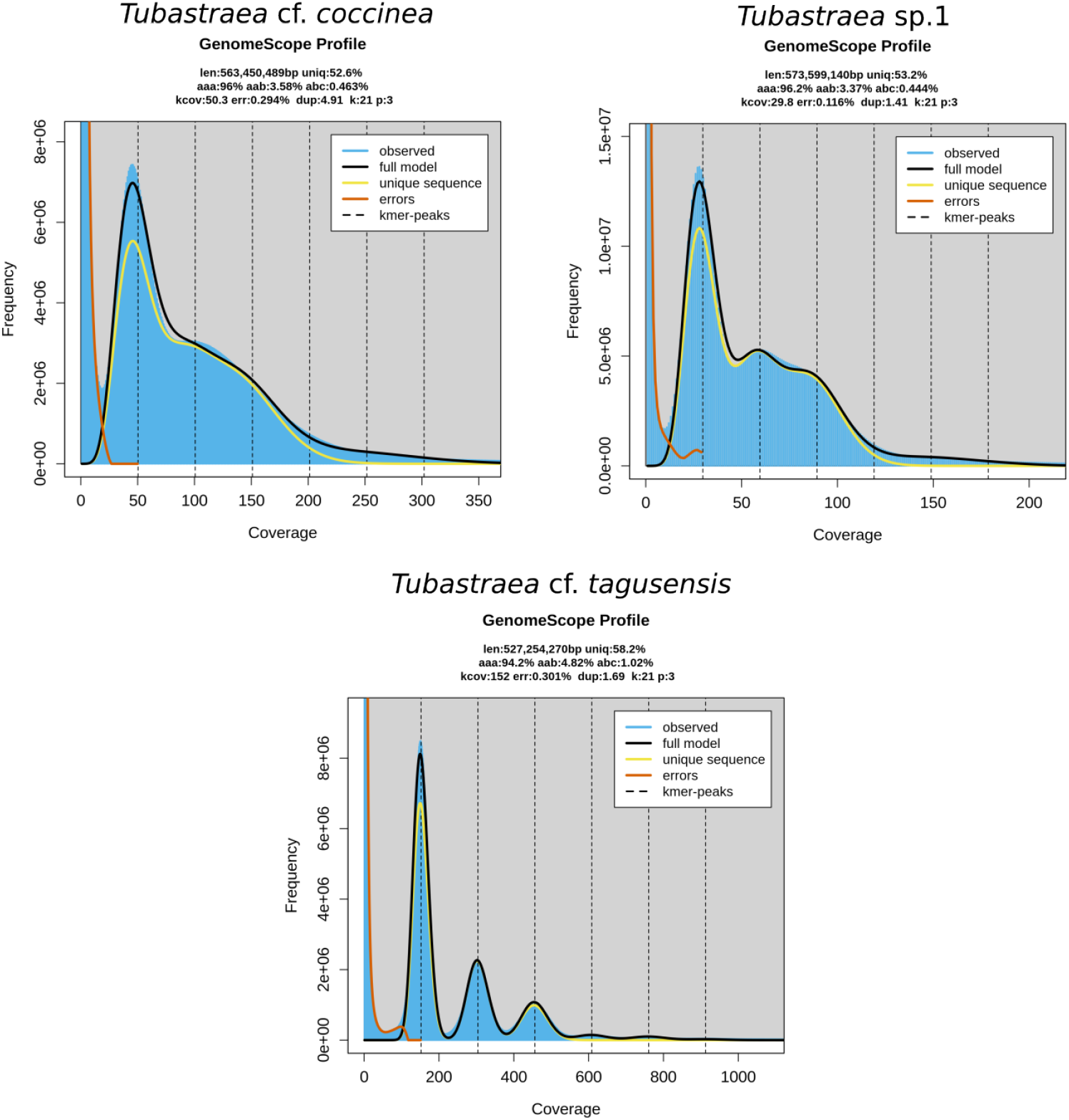
Genomescope linear plots built using Illumina reads from the three morphotypes and kmer size=21bp.

**Figure 8.**
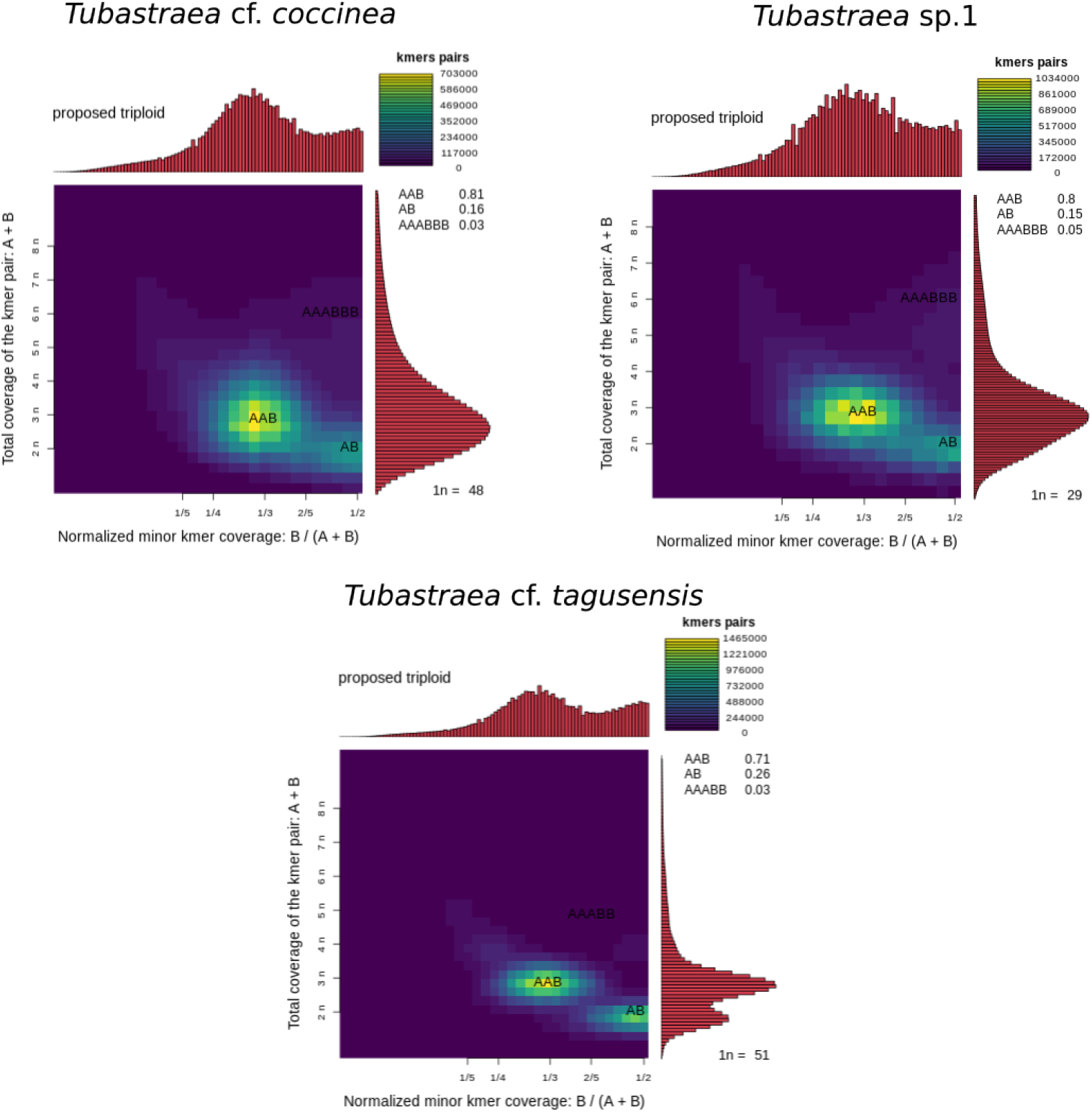
Smudgeplot analysis of *T*. cf. *coccinea, T*. sp. 1, and *T*. cf. *tagusensis*. The heat of each smudge indicates how frequently the haplotype structure is represented in the genome.

**Figure 9.**
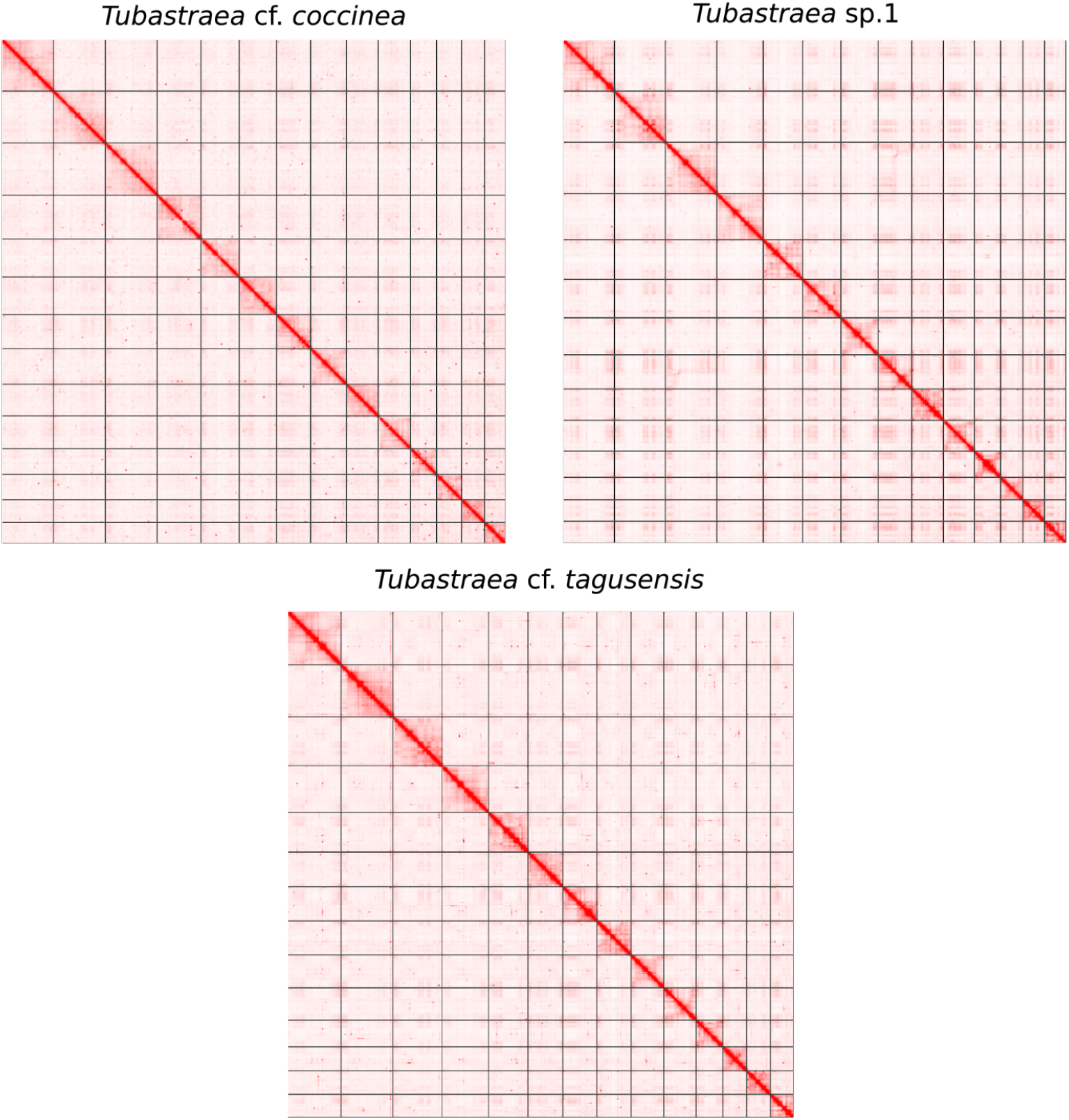
HiC heatmaps of the *T*. cf. *coccinea* and *Tubastraea* sp. 1 morphotype assemblies. Only the 14 chromosome-level scaffolds (pseudochromosomes) are represented.

**Figure 10.**
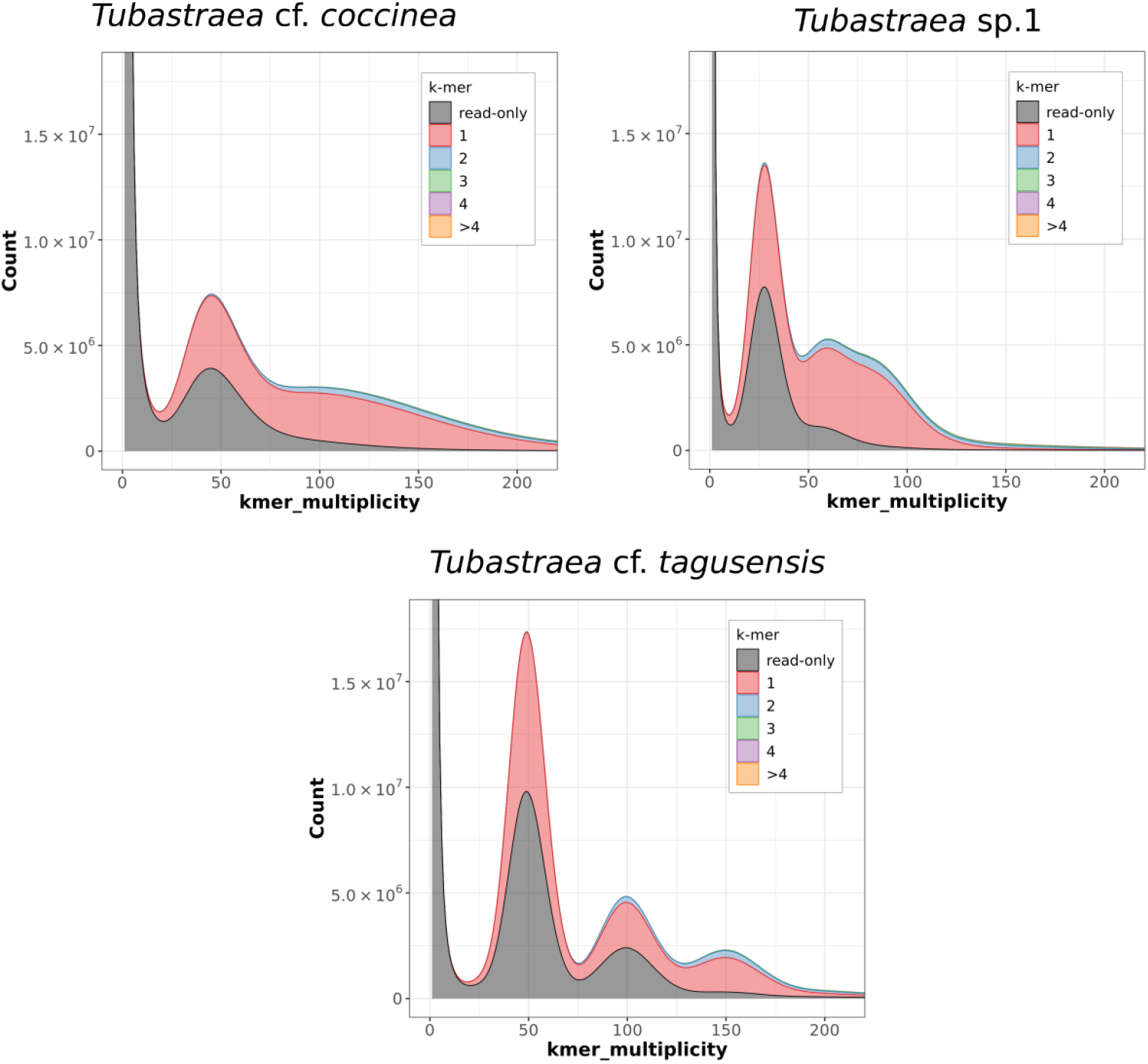
Merqury plots (built using Illumina reads and kmer=21bp) for the three curated morphotype assemblies. Colors represent the number of times the kmer was found in the final assembly.

**Figure 11.**
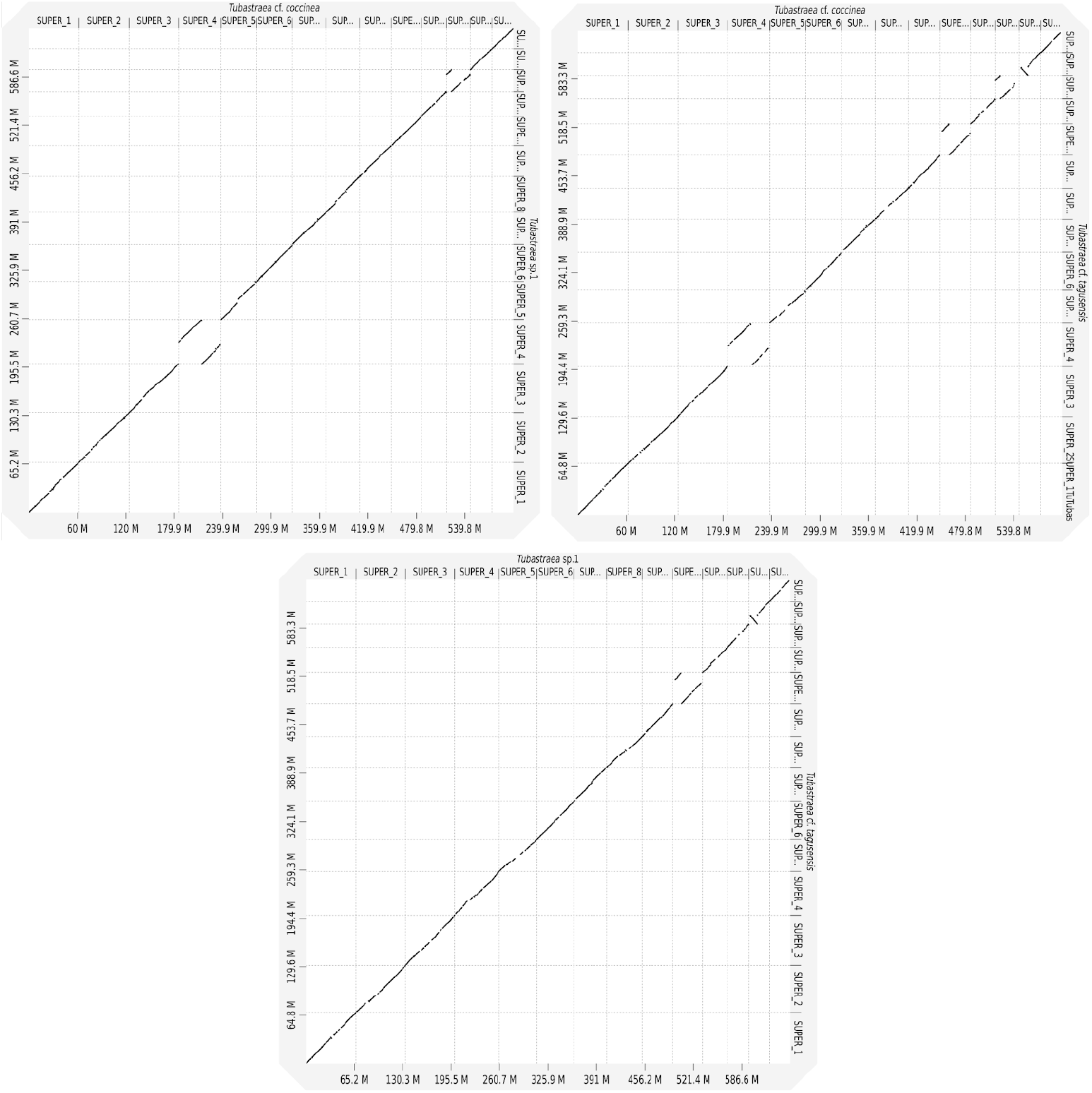
Genome-wide alignment of the nucleotide sequences from each pair of morphotype assemblies. The plots show how the 14 chromosome-level scaffolds of each genome correspond to one another.

**Figure 12.**
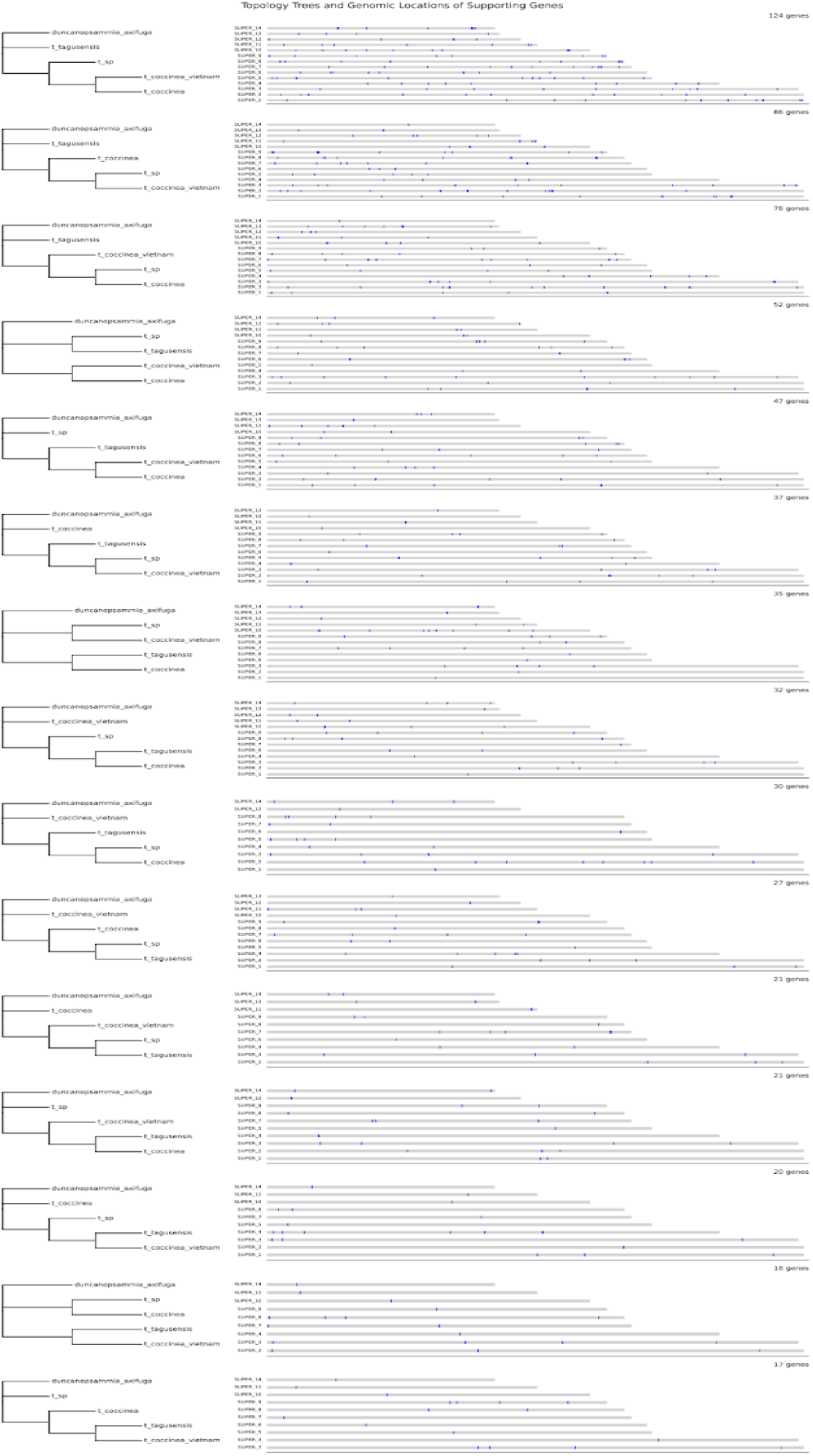
Alternative gene tree topologies and genomic distribution of supporting genes. Left: the 15 topologies recovered across individual gene trees. Right: chromosomal positions of genes supporting each topology, mapped onto the *T*. cf. *coccinea* reference genome. Each horizontal bar represents a chromosome-level scaffold, with blue marks indicating the location of genes supporting the corresponding topology.

**Figure 13.**
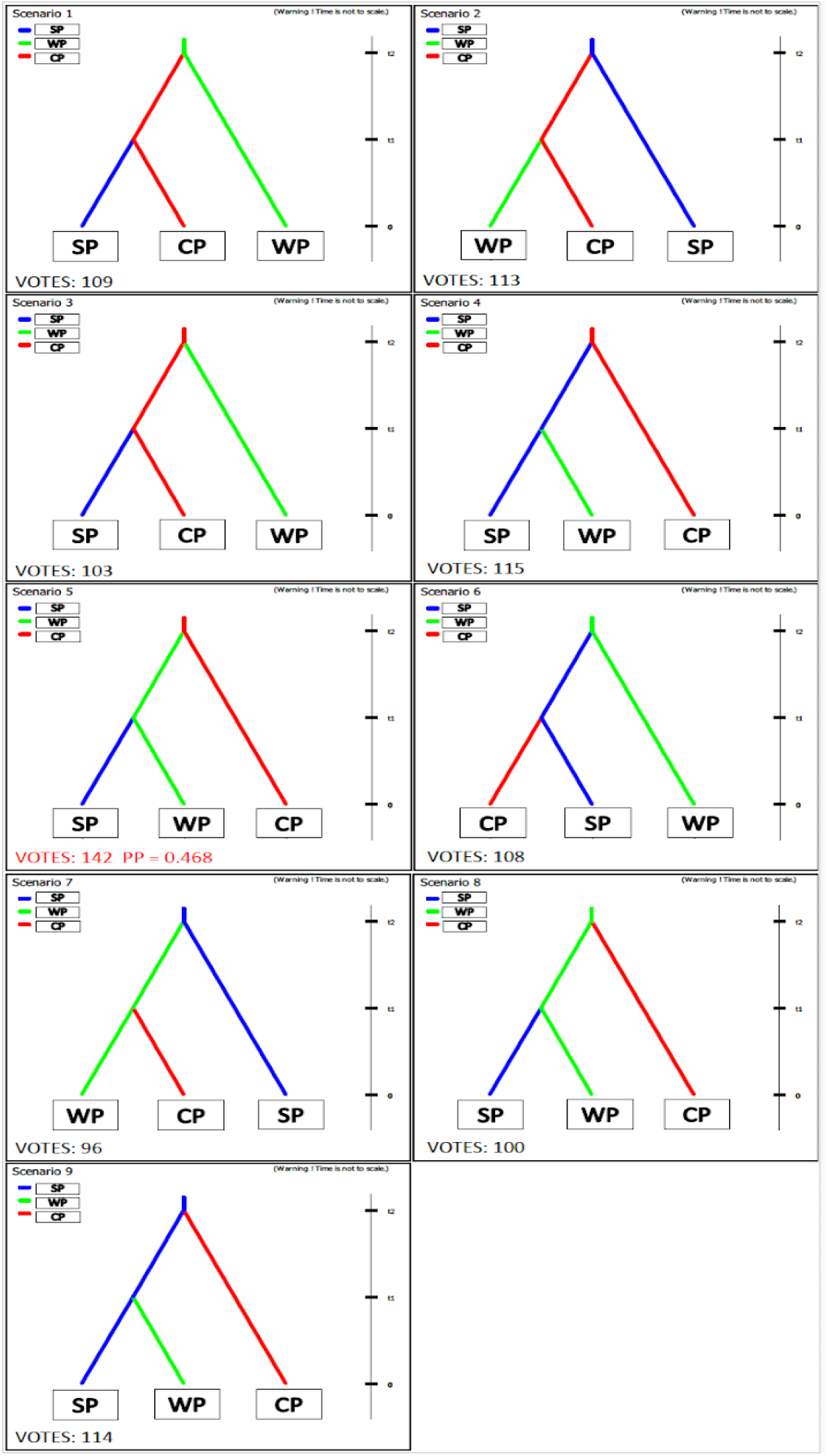
Graphical representation of ABC scenarios to infer the routes of *T. coccinea*, first analysis. Pop 1/N1 = Southeastern Pacific (SP), Pop2/N2 = Western Pacific (WP) and Pop 3/N3 = Centralwestern Pacific (CP). The number of votes obtained for each scenario is shown in black, and in red is the selected as most likely. Posterior probability associated with the most likely scenario is shown.

**Figure 14.**
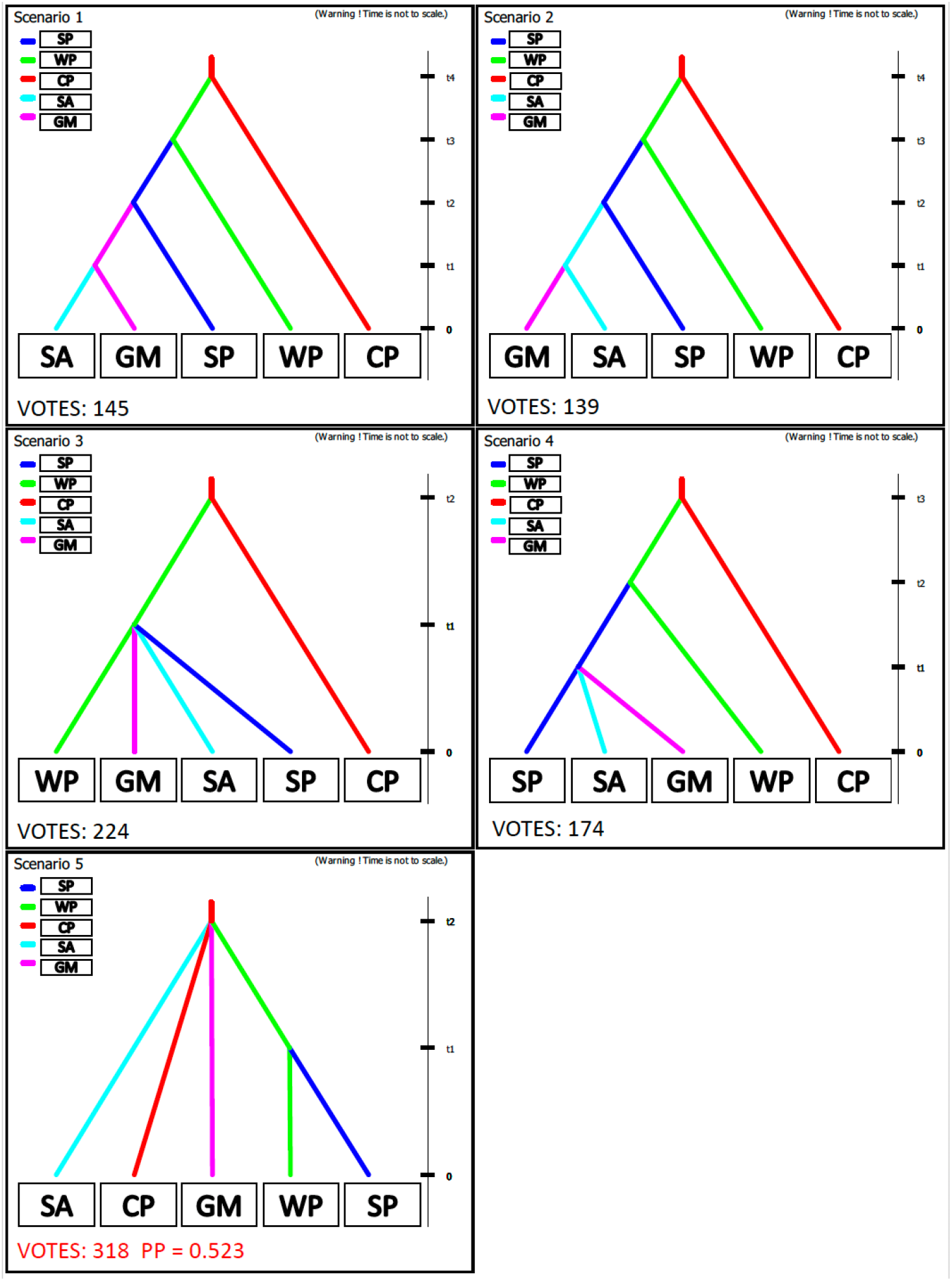
Graphical representation of ABC scenarios to infer the routes of *T. coccinea*, second analysis. Pop 1/N1 = Southeastern Pacific (SA), Pop2/N2 = Western Pacific (WP), Pop 3/N3 = Centralwestern Pacific (CP), Pop 4/N4 = Southwestern Atlantic (SA) and Pop 5/N5 = Gulf of Mexico (GM). The number of votes obtained for each scenario is shown in black, and in red is the selected as most likely. Posterior probability associated with the most likely scenario is shown.

## Supplementary Data

### F. Detailed Modifications to the CTAB-based DNA Extraction Protocol

Soft tissue from sun coral specimens retrieved from aquaria were rinsed with distilled water and immersed in 1.0 ml of CTAB buffer [2% (m/v) CTAB (Sigma-Aldrich), 1.4 M NaCl, 20 mM EDTA, 100 mM Tris-HCl (pH8,0)], with 10 µg of proteinase K (Invitrogen) and 2% of 2-mercaptoethanol (Sigma-Aldrich), freshly added, per 100 mg of tissue. Then, the tissue was kept in a lysis buffer for 4 days, with occasional inversion to promote tissue lysis. Tubes with tissue and buffer were then exposed to freezing with liquid nitrogen for 30 seconds and then thawed and heated to 65ºC in a heat block for approximately 3 minutes. Three freeze-thaw cycles were performed. Then to remove protein and lipids one wash was performed with Phenol:Chloroform:Isoamyl Alcohol (25:24:1) (Sigma-Aldrich), and two washes with Chloroform:Isoamyl Alcohol (24:1) (Sigma-Aldrich). The supernatant was transferred to a new tube containing 1 ml of C4 solution from a Power Soil DNA Isolation kit (MO BIO Laboratories), homogenized by inversion and loaded in a spin column from the DNeasy Blood & Tissue kit (Qiagen, Germany). A final wash was performed with 500 µl of C5 solution. DNA elution was done with 150 µl of Tris-HCl buffer (10 mM, pH 8.5) in three sequencing centrifugation steps.

## Bibliography

Allen Coral Atlas. allencoralatlas.org, (2022). URL https://allencoralatlas.org/.

Arrigoni, R., Kitano, Y. F., Stolarski, J., Hoeksema, B. W., Fukami, H., Stefani, F., Galli, P., Montano, S., Castoldi, E., and Benzoni, F. (2014). A phylogeny reconstruction of the Dendrophylliidae (Cnidaria, Scleractinia) based on molecular and micromorphological criteria, and its ecological implications. Zoologica Scripta, 43(6):661–688. doi: 10.1111/zsc.12072.

Azmi, F., Hewitt, C. L., and Campbell, M. L. (2015). A hub and spoke network model to analyse the secondary dispersal of introduced marine species in Indonesia. ICES Journal of Marine Science, 72(3):1069–1077. doi: 10.1093/icesjms/fsu150.

Bastos, N., Calazans, S. H., Altvater, L., Neves, E. G., Trujillo, A. L., Sharp, W. C., Hoffman, E. A., and Coutinho, R. (2022). Western Atlantic invasion of sun corals: incongruence between morphology and genetic delimitation among morphotypes in the genus Tubastraea. Bulletin of Marine Science. doi: 10.5343/bms.2021.0031.

Bastos, N., Poubel Tunala, L., and Coutinho, R. (2024). Life history strategy of Tubastraea spp. corals in an upwelling area on the Southwest Atlantic: growth, fecundity, settlement, and recruitment. PeerJ, 12:e17829. doi: 10.7717/peerj.17829.

Bastos, N. B. Identificação de espécies, biologia reprodutiva e dinâmica populacional de corais do gênero Tubastraea no sudeste do Brasil. PhD. dissertation, UNIVERSIDADE FEDERAL FLUMINENSE, Programa de Pós-graduação em Dinâmica dos oceanos e da Terra, (2020).

Boavida, J., Ronan Becheler, Choquet, M., Frank, N., Taviani, M., Bourillet, J., Meistertzheim, A., Grehan, A., Savini, A., and Arnaud-Haond, S. (2019). Out of the Mediterranean? Post-glacial colonization pathways varied among cold-water coral species. Journal of Biogeography, 46(5):915–931. doi: 10.1111/jbi.13570.

Borowiec, M. L. (2016). AMAS: a fast tool for alignment manipulation and computing of summary statistics. PeerJ, 4:e1660. doi: 10.7717/peerj.1660. Publisher: PeerJ Inc.

Brown, J. M. and Lemmon, A. R. (2007). The Importance of Data Partitioning and the Utility of Bayes Factors in Bayesian Phylogenetics. Systematic Biology, 56(4):643–655. doi: 10.1080/10635150701546249.

Cairns, S. D. (1994). Scleractinia of the temperate North Pacific. Smithsonian Contributions to Zoology, (557):i–150. doi: 10.5479/si.00810282.557.i.

Cairns, S. D. (2000). A revision of the shallow-water azooxanthellate Scleractinia of the western Atlantic. Studies on the Natural History of the Caribbean Region. Smithsonian Institution, 75:240.

Cairns, S. D. (2001). A generic revision and phylogenetic analysis of the Dendrophylliidae (Cnidaria: Scleractinia). Smithsonian Contributions to Zoology, (615):1–75. doi: 10.5479/si.00810282.615.

Capel, K., Migotto, A., Zilberberg, C., Lin, M., Forsman, Z., Miller, D., and Kitahara, M. (2016). Complete mitochondrial genome sequences of Atlantic representatives of the invasive Pacific coral species Tubastraea coccinea and T. tagusensis (Scleractinia, Dendrophylliidae): Implications for species identification. Gene, 590(2):270–277. doi: 10.1016/j.gene.2016.05.034.

Capel, K. C. C., Toonen, R. J., Rachid, C. T., Creed, J. C., Kitahara, M. V., Forsman, Z., and Zilberberg, C. (2017). Clone wars: asexual reproduction dominates in the invasive range of Tubastraea spp. (Anthozoa: Scleractinia) in the South-Atlantic Ocean. PeerJ, 5:e3873. doi: 10.7717/peerj.3873.

Capel, K. C. C., Zilberberg, C., and Kitahara, D. M. Sistemática do gênero Tubastraea (Scleractinia: Dendrophylliidae) e estrutura genética das espécies invasoras do Atlântico Sul Ocidental. PhD thesis, Universidade Federal do Rio de Janeiro, Rio de Janeiro, (2018).

Capel, K. C. C., Creed, J., Kitahara, M. V., Chen, C. A., and Zilberberg, C. (2019). Multiple introductions and secondary dispersion of Tubastraea spp. in the Southwestern Atlantic. Scientific Reports, 9(1):13978. doi: 10.1038/s41598-019-50442-3.

Carlton, J. T. and Schwindt, E. (2024). The assessment of marine bioinvasion diversity and history. Biological Invasions, 26(1):237–298. doi: 10.1007/s10530-023-03172-7.

Carpenter, K. E., Harrison, P. L., Hodgson, G., Alsaffar, A. H., and Alhazeem, S. H. The Corals and Coral Reef Fishes of Kuwait. Number 7 in Biological Sciences Faculty Books. (1997). URL https://digitalcommons.odu.edu/biology_books/7.

Carrasco, L. A., Doroshov, S., Penman, D. J., and Bromage, N. (1998). Long-term, quantitative analysis of gametogenesis in autotriploid rainbow trout, Oncorhynchus mykiss. Journal of Reproduction and Fertility, 113(2):197–210. doi: 10.1530/jrf.0.1130197.

Chen, X., Han, W., Chang, X., Tang, C., Chen, K., Bao, L., Zhang, L., Hu, J., Wang, S., and Bao, Z. (2025). High-quality genome assembly of the azooxanthellate coral Tubastraea coccinea (Lesson, 1829). Scientific Data, 12(1):507. doi: 10.1038/s41597-02504839-7. Publisher: Nature Publishing Group.

Cornuet, J.-M., Pudlo, P., Veyssier, J., Dehne-Garcia, A., Gautier, M., Leblois, R., Marin, J.-M., and Estoup, A. (2014). DIYABC v2.0: a software to make approximate Bayesian computation inferences about population history using single nucleotide polymorphism, DNA sequence and microsatellite data. Bioinformatics, 30(8):1187–1189. doi: 10.1093/bioinformatics/btt763.

Costantini, M., Guida, F., Amorim, C. G., Da Nóbrega, L. B., Esposito, R., Zupo, V., and Fleury, B. G. (2025). Isolation and Identification of Inter-Correlated Genes from the Invasive Sun Corals Tubastraea Coccinea and Tubastraea Tagusensis (Scleractinia, Cnidaria). International Journal of Molecular Sciences, 26(15):7235. doi: 10.3390/ijms26157235.

Couto, T. D. T. C., Omena, E. P., Oigman-Pszczol, S. S., and Junqueira, A. O. R. (2021). A Method to Assess the Risk of Sun Coral Invasion in Marine Protected Areas. Anais da Academia Brasileira de Ciências, 93:e20200583. doi: 10.1590/0001-3765202120200583. Publisher: Academia Brasileira de Ciências.

Creed, J. C. (2016). Two invasive alien azooxanthellate corals, Tubastraea coccinea and Tubastraea tagusensis, dominate the native zooxanthellate Mussismilia hispida in Brazil. Coral Reefs, 25(3):350–350. doi: 10.1007/s00338-006-0105-x.

Dawson, W., Moser, D., Van Kleunen, M., Kreft, H., Pergl, J., Pyšek, P., Weigelt, P., Winter, M., Lenzner, B., Blackburn, T. M., Dyer, E. E., Cassey, P., Scrivens, S. L., Economo, E. P., Guénard, B., Capinha, C., Seebens, H., García-Díaz, P., Nentwig, W., García-Berthou, E., Casal, C., Mandrak, N. E., Fuller, P., Meyer, C., and Essl, F. (2017). Global hotspots and correlates of alien species richness across taxonomic groups. Nature Ecology & Evolution, 1(7):0186. doi: 10.1038/s41559-017-0186.

de Oliveira Soares, M., Davis, M., de Paiva, C. C., and de Macêdo Carneiro, P.B. (2016). Mesophotic ecosystems: coral and fish assemblages in a tropical marginal reef (northeastern Brazil). Marine Biodiversity, 48(3):1631–1636. doi: 10.1007/s12526-0160615-x.

de Paula, A. F. and Creed, J. C. (2004). Two species of the coral Tubastraea (Cnidaria, Scleractinia) in Brazil: a case of accidental introduction.

De Paula, A. F., De Oliveira Pires, D., and Creed, J. C. (2014). Reproductive strategies of two invasive sun corals (Tubastraea spp.) in the southwestern Atlantic. Journal of the Marine Biological Association of the United Kingdom, 94(3):481–492. doi: 10.1017/S0025315413001446.

Dutra, B. S. V. M., Carlos-Júnior, L. A., and Creed, J. C. (2023). When species become invasive research becomes problem oriented: a synthesis of knowledge of the stony coral Tubastraea. Biological Invasions, 25(7):2069–2088. doi: 10.1007/s10530-02303032-4.

Ehrenberg. Dritter Beitrag zur Erkenntniss grosser Organisation in der Richtung des kleinsten Raumes. Abhandlungen der Königlichen Akademie der Wissenschaften zu Berlin, volume 1834. Realschul-Buchhandlung, Berlin, (1834). URL https://www.biodiversitylibrary.org/item/93981. ISSN: 2944-4411 Pages: 1-1002.

Felsenstein, J. (1985). Confidence limits on phylogenies: an approach using the bootstrap. Evolution; International Journal of Organic Evolution, 39(4):783–791. doi: 10.1111/j.1558-5646.1985.tb00420.x.

Fenner, D. (1999). New observations on the stony coral (Scleractinia, Milleporidae, and Stylasteridae) species of Belize (Central America) and Cozumel (Mexico). ResearchGate.

Fenner, D. and Banks, K. (2004). Orange Cup Coral Tubastraea coccinea invades Florida and the Flower Garden Banks, Northwestern Gulf of Mexico. Coral Reefs. doi: 10.1007/s00338-004-0422-x.

Figueroa, D. F., McClure, A., Figueroa, N. J., and Hicks, D. W. (2019). Hiding in plain sight: invasive coral <I>Tubastraea tagusensis</i> (Scleractinia:Hexacorallia) in the Gulf of Mexico. Coral Reefs, 38(3):395–403. doi: 10.1007/s00338-019-01807-7.

Fraimout, A., Debat, V., Fellous, S., Hufbauer, R. A., Foucaud, J., Pudlo, P., Marin, J.-M., Price, D. K., Cattel, J., Chen, X., Deprá, M., François Duyck, P., Guedot, C., Kenis, M., Kimura, M. T., Loeb, G., Loiseau, A., Martinez-Sañudo, I., Pascual, M., Polihronakis Richmond, M., Shearer, P., Singh, N., Tamura, K., Xuéreb, A., Zhang, J., and Estoup, A. (2017). Deciphering the routes of invasion of Drosophila suzukii by means of ABC random forest. Molecular Biology and Evolution, page msx050. doi: 10.1093/molbev/msx050.

Garcia, G. D., Gregoracci, G. B., Santos, E. d. O., Meirelles, P. M., Silva, G. G. Z., Edwards, R., Sawabe, T., Gotoh, K., Nakamura, S., Iida, T., de Moura, R. L., and Thompson, F. L. (2013). Metagenomic analysis of healthy and white plague-affected Mussismilia braziliensis corals. Microbial Ecology, 65(4):1076–1086. doi: 10.1007/s00248-0120161-4.

Hodgson, G. and Carpenter, K. (1995). Scleractinian Corals of Kuwait!

Hoeksema, B. W., Samimi-Namin, K., and Vermeij, M. J. A. (2024). Neutral interactions among three nonindigenous coral species in a tropical marine fouling community. Ecology, 105(8):e4371. doi: 10.1002/ecy.4371. _eprint: https://esajournals.onlinelibrary.wiley.com/doi/pdf/10.1002/ecy.4371.

Holmes, S. (2003). Bootstrapping Phylogenetic Trees: Theory and Methods. Statistical Science, 18(2). doi: 10.1214/ss/1063994979.

Howe, K., Chow, W., Collins, J., Pelan, S., Pointon, D.-L., Sims, Y., Torrance, J., Tracey, A., and Wood, J. (2021). Significantly improving the quality of genome assemblies through curation. GigaScience, 10(1):giaa153. doi: 10.1093/gigascience/giaa153.

Kainer, D. and Lanfear, R. (2015). The Effects of Partitioning on Phylogenetic Inference. Molecular Biology and Evolution, 32(6):1611–1627. doi: 10.1093/molbev/msv026.

Kalyaanamoorthy, S., Minh, B. Q., Wong, T. K. F., von Haeseler, A., and Jermiin, L. S. (2017). ModelFinder: fast model selection for accurate phylogenetic estimates. Nature Methods, 14(6):587–589. doi: 10.1038/nmeth.4285. Publisher: Nature Publishing Group.

Katoh, K., Rozewicki, J., and Yamada, K. D. (2019). MAFFT online service: multiple sequence alignment, interactive sequence choice and visualization. Briefings in Bioinformatics, 20(4):1160–1166. doi: 10.1093/bib/bbx108.

Kimura, M. (1980). A simple method for estimating evolutionary rates of base substitutions through comparative studies of nucleotide sequences. Journal of Molecular Evolution, 16 (2):111–120. doi: 10.1007/BF01731581.

Knowlton, N. Sibling Species in the Sea. Annual Review of Ecology and Systematics, volume 24. JSTOR, (1993).

Lages, B., Fleury, B., Menegola, C., and Creed, J. (2011). Change in tropical rocky shore communities due to an alien coral invasion. Marine Ecology Progress Series, 438:85–96. doi: 10.3354/meps09290.

Lesson, R.-P. Voyage autour du monde: exécuté par ordre du roi, sur la corvette de Sa Majesté, la Coquille, pendant les années 1822, 1823, 1824, et 1825, volume t.2:pt.1 (1830) [Zoologie Text]. Arthus Bertrand, Paris, (1830). doi: 10.5962/bhl.title.57936. URL https://www.biodiversitylibrary.org/item/119040. Pages: 1-488.

Li, H. (2018). Minimap2: pairwise alignment for nucleotide sequences. Bioinformatics, 34 (18):3094–3100. doi: 10.1093/bioinformatics/bty191.

Luz, B. L. P., Di Domenico, M., Migotto, A. E., and Kitahara, M. V. (2020). Life-history traits of Tubastraea coccinea: Reproduction, development, and larval competence. Ecology and Evolution, 10(13):6223–6238. doi: 10.1002/ece3.6346.

López, C., Clemente, S., Moreno, S., Ocaña, O., Herrera, R., Moro, L., Monterroso, O., Rodríguez, A., and Brito, A. (2019). Invasive Tubastraea spp. and Oculina patagonica and other introduced scleractinians corals in the Santa Cruz de Tenerife (Canary Islands) harbor: Ecology and potential risks. Regional Studies in Marine Science, 29:100713. doi: 10.1016/j.rsma.2019.100713.

Mantelatto, M. C., Creed, J. C., Mourão, G. G., Migotto, A. E., and Lindner, A. (2011). Range expansion of the invasive corals Tubastraea coccinea and Tubastraea tagusensis in the Southwest Atlantic. Coral Reefs, 30(2):397–397. doi: 10.1007/s00338-011-0720-z.

Marshall, D. C., Simon, C., and Buckley, T. R. (2006). Accurate Branch Length Estimation in Partitioned Bayesian Analyses Requires Accommodation of Among-Partition Rate Variation and Attention to Branch Length Priors. Systematic Biology, 55(6):993–1003. doi: 10.1080/10635150601087641.

Martins, I., Capel, K. C. C., and Abessa, D. M. D. S. (2024). Adults of Sun Coral Tubastraea coccinea(Lesson 1829) Are Resistant to New Antifouling Biocides. Toxics, 12(1):44. doi: 10.3390/toxics12010044.

McFADDEN, C. S., Benayahu, Y., Pante, E., Thoma, J. N., Nevarez, P. A., and France, S. C. (2011). Limitations of mitochondrial gene barcoding in Octocorallia. Molecular Ecology Resources, 11(1):19–31. doi: 10.1111/j.1755-0998.2010.02875.x.

Mera-Rodríguez, D., Fernández-Marín, H., and Rabeling, C. (2025). Phylogenomic approach to integrative taxonomy resolves a century-old taxonomic puzzle and the evolutionary history of the Acromyrmex octospinosus species complex. Systematic Entomology, 50(3):469–494. doi: 10.1111/syen.12665. _eprint:https://resjournals.onlinelibrary.wiley.com/doi/pdf/10.1111/syen.12665.

Minh, B. Q., Schmidt, H. A., Chernomor, O., Schrempf, D., Woodhams, M. D., von Haeseler, A., and Lanfear, R. (2020). IQ-TREE 2: New Models and Efficient Methods for Phylogenetic Inference in the Genomic Era. Molecular Biology and Evolution, 37(5):1530–1534. doi: 10.1093/molbev/msaa015.

Nascimento, N. F. et al. (2017). Growth, fatty acid composition, and reproductive parameters of diploid and triploid yellowtail tetra Astyanax altiparanae. Aquaculture, 471:163–171.

Neves da Rocha, L.S., Nunes, J. A. C. C., Miranda, R. J., and Kikuchi, R. K. P. (2024). Effects of invasive sun corals on habitat structural complexity mediate reef trophic pathways. Marine Biology, 171(4):76. doi: 10.1007/s00227-024-04394-6.

of the Environment – MMA, M., Brazilian Institute of the Environment and Renewable Natural Resources – Ibama, and Chico Mendes Institute for Biodiversity Conservation – ICMBio. Diagnóstico sobre a invasão do coral-sol (Tubastraea spp.) no Brasil. Technical report, (2018). URL https://www.ibama.gov.br/phocadownload/consultapublica/2018/2018-01-diagnostico-coral-sol-consulta-publica_revisaoMMA.pdf?utm_source=chatgpt.com.

Precht, W. F., Hickerson, E. L., Schmahl, G. P., and Aronson, R. B. (2014). The Invasive Coral Tubastraea coccinea (Lesson, 1829): Implications for Natural Habitats in the Gulf of Mexico and the Florida Keys. Gulf of Mexico Science, 32(1). doi: 10.18785/goms.3201.05.

Puillandre, N., Brouillet, S., and Achaz, G. (2020). ASAP: assemble species by automatic partitioning. Molecular Ecology Resources, 21(2):609–620. doi: 10.1111/1755-0998.13281.

Quattrini, A. M., Snyder, K. E., Purow-Ruderman, R., Seiblitz, I. G. L., Hoang, J., Floerke, N., Ramos, N. I., Wirshing, H. H., Rodriguez, E., and McFadden, C. S. (2023). Mitonuclear discordance within Anthozoa, with notes on unique properties of their mitochondrial genomes. Scientific Reports, 13(1):7443. doi: 10.1038/s41598-023-34059-1.

Ranallo-Benavidez, T. R., Jaron, K. S., and Schatz, M. C. (2020). GenomeScope 2.0 and Smudgeplot for reference-free profiling of polyploid genomes. Nature Communications, 11(1):1432. doi: 10.1038/s41467-020-14998-3. Publisher: Nature Publishing Group.

Razak, T. B., Boström-Einarsson, L., Alisa, C. A. G., Vida, R. T., and Lamont, T. A. (2022). Coral reef restoration in Indonesia: a review of policies and projects. Marine Policy, 137: 104940. doi: 10.1016/j.marpol.2021.104940.

Rhie, A., Walenz, B. P., Koren, S., and Phillippy, A. M. (2020). Merqury: reference-free quality, completeness, and phasing assessment for genome assemblies. Genome Biology, 21 (1):245. doi: 10.1186/s13059-020-02134-9.

Romano, S. L. and Cairns, S. D. (2000). Molecular Phylogenetic Hypotheses for The Evolution of Scleractinian Corals. Bulletin of Marine Science, 67:1043–1068.

Sammarco, P., Porter, S., and Cairns, S. (2010). A new coral species introduced into the Atlantic Ocean - Tubastraea micranthus (Ehrenberg 1834) (Cnidaria, Anthozoa, Scleractinia): An invasive threat? Aquatic Invasions, 5(2):131–140. doi: 10.3391/ai.2010.5.2.02.

Sammarco, P., Atchison, A., and Boland, G. (2014). Coral Settlement on Oil/Gas Platforms in the Northern Gulf of Mexico: Preliminary Evidence of Rarity. Gulf of Mexico Science, 32(1). doi: 10.18785/goms.3201.02.

Sammarco, P., Porter, S., Sinclair, J., and Genazzio, M. (2014). Population expansion of a new invasive coral species, Tubastraea micranthus, in the northern Gulf of Mexico. Marine Ecology Progress Series, 495:161–173. doi: 10.3354/meps10576.

Sammarco, P. W., Brazeau, D. A., and Sinclair, J. (2012). Genetic Connectivity in Scleractinian Corals across the Northern Gulf of Mexico: Oil/Gas Platforms, and Relationship to the Flower Garden Banks. PLoS ONE, 7(4):e30144. doi: 10.1371/journal.pone.0030144.

Sampaio, C. L. S., Miranda, R. J., Maia-Nogueira, R., and Nunes, J. d. A. C. C. (2012). New occurrences of the nonindigenous orange cup corals Tubastraea coccinea and T. tagusensis (Scleractinia: Dendrophylliidae) in Southwestern Atlantic. Check List, 8(3):528–530. doi: 10.15560/8.3.528. Publisher: Pensoft Publishers.

Searcy, C. A., Howell, H. J., David, A. S., Rumelt, R. B., and Clements, S. L. (2023). Patterns of Non-Native Species Introduction, Spread, and Ecological Impact in South Florida, the World’s Most Invaded Continental Ecoregion. Annual Review of Ecology, Evolution, and Systematics, 54(1):195–218. doi: 10.1146/annurev-ecolsys-110421-103104.

Seiblitz, I. G. L., Vaga, C. F., Capel, K. C. C., Cairns, S. D., Stolarski, J., Quattrini, A. M., and Kitahara, M. V. (2022). Caryophylliids (Anthozoa, Scleractinia) and mitochondrial gene order: Insights from mitochondrial and nuclear phylogenomics. Molecular Phylogenetics and Evolution, 175:107565. doi: 10.1016/j.ympev.2022.107565.

Serra, Bastos, Coutinho, SCHIZAS, JOHNSSON, and Neves. (2024). Four new species of Tubastraea (Scleractinia, Dendrophylliidae) from the Brazilian Coast, Southwestern Atlantic. Instituto de Ecologia y Ciencias Ambientales, 19(2). doi: 10.54451/PanamJAS.19.2.113.

Shearer, T. L., Van Oppen, M. J. H., Romano, S. L., and Wörheide, G. (2002). Slow mitochondrial DNA sequence evolution in the Anthozoa (Cnidaria). Molecular Ecology, 11 (12):2475–2487. doi: 10.1046/j.1365-294X.2002.01652.x.

Simão, F. A., Waterhouse, R. M., Ioannidis, P., Kriventseva, E. V., and Zdobnov, E. M. (2015). BUSCO: assessing genome assembly and annotation completeness with single-copy orthologs. Bioinformatics, 31(19):3210–3212. doi: 10.1093/bioinformatics/btv351.

Soares, M. O., Pereira, P. H., Feitosa, C. V., Maggioni, R., Rocha, R. S., Bezerra, L. E. A., Duarte, O. S., Paiva, S. V., Noleto-Filho, E., Silva, M. Q. M., Csapo-Thomaz, M., Garcia, T. M., Arruda Júnior, J. P. V., Cottens, K. F., Vinicius, B., Araújo, R., Eirado, C. B. D., Santos, L. P. S., Guimarães, T. C. S., Targino, C. H., Amorim-Reis Filho, J., Santos, W. C. R. D., Klautau, A. G. C. D. M., Gurjão, L. M. D., Machado, D. A. N., Maia, R. C., Santos, E. S., Sabry, R., Asp, N., Carneiro, P. B., Rabelo, E. F., Tavares, T. C., Lima, G. V. D., Sampaio, C. L., Rocha, L. A., Ferreira, C. E., and Giarrizzo, T. (2023). Lessons from the invasion front: Integration of research and management of the lionfish invasion in Brazil. Journal of Environmental Management, 340:117954. doi: 10.1016/j.jenvman.2023.117954.

Souza Filho, J. R. S. Cruz, I.C., Martinez, S. T., and De Andrade, J. B. (2024). Bioinvasion of Sun Coral (Tubastraea tagusensis): A Threat to the Essential Ecosystem Services in Morro de São Paulo, Cairú, Bahia, Brazil. Journal of Coastal Research, 113(Sp1). doi: 10.2112/JCR-SI113-029.1.

Stephens, T. G., Strand, E. L., Putnam, H. M., and Bhattacharya, D. (2023). Ploidy Variation and Its Implications for Reproduction and Population Dynamics in Two Sympatric Hawaiian Coral Species. Genome Biology and Evolution, 15(8):evad149. doi: 10.1093/gbe/evad149.

Subhan, B., Razak, T. B., Arafat, D., Zamani, N. P., Prehadi Lestari, D.F., and Hoeksema, B. W. (2022). A New Northernmost Distribution Record of the Reef Coral Duncanopsammia axifuga at Bird’s Head Peninsula, West Papua, Indonesia. Diversity, 14(9):713. doi: 10.3390/d14090713. Publisher: Multidisciplinary Digital Publishing Institute.

Tabatabaee, Y., Zhang, C., Warnow, T., and Mirarab, S. (2023). Phylogenomic branch length estimation using quartets. Bioinformatics, 39(Supplement_1):i185–i193. doi: 10.1093/bioinformatics/btad221.

Takata, K., Nonaka, M., Xia, F., Kikuchi, T., Gibu, K., and Yasuda, N. (2025). The complete mitochondrial genome of Pleurocorallium inutile (Octocorallia: Scleralcyonacea: Coralliidae). Molecular Biology Reports, 52(1):531. doi: 10.1007/s11033-025-10583-3.

Tamura, K., Stecher, G., and Kumar, S. (2021). MEGA11: Molecular Evolutionary Genetics Analysis Version 11. Molecular Biology and Evolution, 38(7):3022–3027. doi: 10.1093/molbev/msab120.

Tang, H., Krishnakumar, V., Zeng, X., Xu, Z., Taranto, A., Lomas, J. S., Zhang, Y., Huang, Y., Wang, Y., Yim, W. C., Zhang, J., and Zhang, X. (2024). JCVI: A versatile toolkit for comparative genomics analysis. iMeta, 3(4):e211. doi: 10.1002/imt2.211.

Ugarte, P. D. d. S. and Garraffoni, A. R. S. (2024). Removal of historical taxonomic bias and its impact on biogeographic analyses: a case study of Neotropical tardigrade fauna. Zoological Journal of the Linnean Society, 201(3):zlae091. doi: 10.1093/zoolinnean/zlae091.

Veron, J., Devantier, L., Turak, E., Green, A., Kininmonth, S., Stafford-Smith, M., and Peter-son, N. The Coral Triangle. In Coral Reefs: An Ecosystem in Transition, pages 47–55. (2011). ISBN 978-94-007-0113-7. doi: 10.1007/978-94-007-0114-4_5. Journal Abbreviation: Coral Reefs: An Ecosystem in Transition.

Vogt, G., Falckenhayn, C., Schrimpf, A., Schmid, K., Hanna, K., Panteleit, J., Helm, M., Schulz, R., and Lyko, F. (2015). The marbled crayfish as a paradigm for saltational speciation by autopolyploidy and parthenogenesis in animals. Biology Open, 4(11):1583– 1594. doi: 10.1242/bio.014241.

Wells, J. W. (1982). Notes on Indo-Pacific Scleractinian Corals. Part 9. 1 New Corals from the Galapagos Islands. Pacific Science, 36(2):211–219.

Yiu and Qiu. (2022). Three New Species of the Sun Coral Genus Tubastraea (Scleractinia: Dendrophylliidae) from Hong Kong, China. Zoological Studies, (61). doi: 10.6620/ZS.2022.61-45.

Yiu, Chung, and Qiu. (2021). A new species of the sun coral genus Tubastraea (Scleractinia: Dendrophylliidae) from Hong Kong. Zootaxa, 5047(1):1–16. doi: 10.11646/zootaxa.5047.1.1.

Zhang, C. and Mirarab, S. (2022). Weighting by Gene Tree Uncertainty Improves Accuracy of Quartet-based Species Trees. Molecular Biology and Evolution, 39(12):msac215. doi: 10.1093/molbev/msac215.

Zimin, A. V., Marçais, G., Puiu, D., Roberts, M., Salzberg, S. L., and Yorke, J. A. (2013). The MaSuRCA genome assembler. Bioinformatics, 29(21):2669–2677. doi: 10.1093/bioinformatics/btt476.

